# Pathologic and phylogenetic characterization of Usutu virus infections in free-ranging wild birds in Switzerland (2015-2020)

**DOI:** 10.1101/2025.05.01.651669

**Authors:** Simone Roberto Rolando Pisano, Michelle Imlau, Ursula Sattler, Francesco Carlo Origgi

## Abstract

Usutu Virus (USUV) is an emerging flavivirus causing fatal disease in free-ranging and captive birds but also infecting mammals and representing a significant zoonotic agent in Europe, making monitoring and investigations of this viral agent crucial both for conservation and public health.

We carried out a retrospective investigation focusing on free-ranging wild birds confirmed to be USUV infected examined at the Institute for Fish and Wildlife Health (FIWI), Switzerland. The pathology and virology findings together with the phylogeographic and environmental data were examined for underlying factors to the dynamics of the USUV infection in Switzerland, between 2015 and 2020, with particular focus to the major peak of 2018. USUV infection was confirmed by reverse transcription-PCR in 67 birds, with common blackbirds (*Turdus merula*, 73%; 49/67) being overrepresented, showing signs of diseases including poor body condition (84%; 56/67) and splenomegaly (75%; 50/67). Histology (n=34) revealed changes in the heart (58%; 19/33), the central nervous system (67%; 22/33) and the liver (62%; 21/34). Of interest, the splenic changes (86%, 25/29) showed different, but likely correlated phenotypes, characterized by the presence of discrete and/or diffuse histiocytic infiltrates in the congested and lymphoid depleted spleen parenchyma, possibly reflecting different stages of the disease. Phylogenetic analysis unambiguously revealed that the recent USUV infections were predominantly caused by strains of the Europe 3 lineage. USUV Africa 3 strains were detected exclusively from southern Switzerland in one location (two blackbirds).

Our investigation revealed intralesional Usutu viral RNA consistent with infection in previously undescribed hosts (*Corvus frugilegus*, *Cygnus olor*) and provided new insights into the pathogenesis and the ecology of the disease. The environmental data suggested the occurrence of unique weather conditions, possibly boosting the mosquito population and favoring an unprecedented increase of cases in 2018. Finally, the increase in genetic diversity of the circulating strains might have contributed to the recorded high mortality in 2018.

**Author Summary:** At the end of the 20^th^ century, Usutu virus (USUV) was associated for the first time with mass mortality in wild black birds in Italy. Shortly after, reports on avian fatalities associated with USUV all over Europe appeared. In 2018 an unprecedented high occurrence of Usutu virus disease (UVD) caused high mortalities in wild birds in Switzerland and other European countries. Here we examined deceased wild birds with UVD with an emphasis on pathological examination and identification of USUV strains. The spleen is one of the main organs affected and we propose a timeline of lesion progression correlating with early, intermediate and late changes. Whereas we mainly identified strains of the European 3 lineage, we also found a hotspot of African 3 lineage strains. This indicates multiple entries of USUV into Switzerland. Although the disease is especially fatal in Strigiformes and Passeriformes, we expanded the spectrum of bird species (mute swans, rooks) naturally infected with the USUV.

Usutu virus is an arthropod-borne virus and transmission occurs mainly via blood feeding vectors such as mosquitoes. We discuss the influence of weather conditions on mosquito population and the potential role of invasive mosquito species, such as the Asian tiger mosquito and other *Aedes* spp., as additional vectors for disease.

## Introduction

Usutu virus (USUV) is an arthropod-borne flavivirus closely related to West Nile virus and part of the Japanese encephalitis virus group [1]. Similarly to other viruses of its group, USUV is mainly transmitted by mosquitos (mainly *Culex* spp. mosquitos) [2,3]. Phylogenetic studies have shown the separation of USUV into distinct lineages, which are assigned according to their putative geographic origin, namely Africa 1, 2 and 3 and Europe 1, 2, 3, 4 and 5 [4–6]. The virus is named after the region of its original discovery in *Culex neavei*, near the Usutu river, Eswatini (formerly Swaziland) in 1959 [7]. The first confirmed cases in continental Europe included a series of deaths among passerines (Order Passeriformes), mostly common blackbirds (*Turdus merula*) in Vienna, Austria in 2001 [8]. Retrospectively, USUV was associated with a blackbird mass mortality in Tuscany, Italy, in 1996 [9]. Shortly after its first report, presence of DNA of USUV was recorded in wild birds in other European countries including Hungary (2003) [10], Switzerland (2006) [11], Spain (2006) [12], Italy (2008) [13], Germany (2011) [14], Czech Republic (2011-2012) [15], Belgium (2014, 2016, 2017-2018) [16–18], France (2015) [19], the Netherlands (2016) [20] and finally the UK and the Luxembourg (2020) [21,22]. Up to 2021, contact between USUV and animals was reported in fifteen EU countries, 11 of them through molecular testing, especially in birds [23].

To date, infections with USUV have been reported in at least 93 different bird species belonging to 35 families [24]. Affected species belong mainly to Order Passeriformes, and to a minor extent Accipitriformes, Strigiformes, Anseriformes and Galliformes [24]. In Germany, apparent prevalence of USUV in wild birds ranged between 1.2 to 12.2% [25].

Reports of USUV infections in mammals [23] include bats [26], dogs [27], horses [27,28], red deer (*Cervus elaphus*) [29] and humans [30–32]. The range of mammal species showing antibody response to USUV is even broader [23].

The extent and severity of the outbreaks ranges from single deaths in both captive and free-ranging individuals [33] to massive mortalities in the wild [13]. Clinical signs in USUV infected birds are primarily neurological (depression, incoordination, seizures) and peracute death [11]. These signs are associated with gross findings including spleno- and hepatomegaly, multisystemic necrosis (spleen, liver and kidneys) along with encephalitis and myocarditis detected under light microscopy [10,11,20,34,35].

In humans, especially in immunosuppressed individuals, USUV infection can cause encephalitis and meningoencephalitis [36–39], whereas in more endemic regions, such as Africa, a clinically silent viremia or the so called *Usutu fever*, a mild disease characterized by fever, rash, jaundice and headache [40], is the most common outcome of the infection. Widespread epidemiologic data on prevalence in African countries is lacking. Studies on seroprevalence in Europe, including Italy, Germany and Serbia, documented USUV-antibody prevalence between 0.02-1.1% among healthy blood donors, consistent with a non-negligible risk of USUV infections in humans [41].

The Institute for Fish and Wildlife Health (FIWI) at the University of Bern regularly receive carcasses along with tissue samples of wild animals from all the 26 Swiss cantons (administrative units) and additionally from the neighboring Principality of Liechtenstein in the framework of a national surveillance program for wildlife health [42]. Following the first detection of USUV around the city of Zurich, Switzerland in 2006 [11], cases have been sporadically observed by the FIWI during the following years but more regularly since 2015 (unpublished data). Interestingly, a sudden and major increase of cases occurred in late summer and through the fall of 2018, paralleling what was recorded in other European countries, including Austria [43], Hungary [43], Italy [31], Croatia [44], Slovakia [45], Germany [46], Belgium [17] and the Netherlands [47], consistent with an epidemic sweeping through the European continent. This similar and unusual pan-European epidemiological picture, with a strikingly overlapping timing corresponding to the late summer and fall 2018 prompt us to investigate the features of the Swiss epidemic as a putative “proxy” of the European scenario. In particular, we carried out a thorough investigation of the associated pathology and of the infecting viral strains together with the available environmental data in order to determine if any of these aspects or their combination, might provide an explanation of the dynamic of the 2018 epidemic.

Here we report the results of our investigation, (1) providing new pathology data, consistent with a putative involvement of histiocytes in the pathogenesis of USUV-associated disease; (2) showing that the same USUV lineage (Europe 3) was circulating during, prior and after 2018, making unlikely the emergence of a highly virulent virus responsible of the 2018 epidemic. However, a relevant genetic diversity was seen within the circulating strains in 2018 and a lineage not detected prior was recorded in Switzerland. (3) Finally, circumstantial evidence of particularly favorable environmental conditions occurring in 2018 might have boosted the vector population leading eventually to an increase in fatalities.

## Results

### Animals

USUV positivity was obtained for sixty-seven birds by RT-PCR and it was supported by successful and readable Sanger sequencing for 49 of them (83%) (Table 3). The USUV positive birds belonged to three orders (64 Passeriformes, 1 Apodiformes and 2 Anseriformes) and included eleven species and more specifically, 73% (49/67) common blackbirds (*Turdus merula*), 7% (5/67) rooks (*Corvus frugilegus*), 4% (3/67) carrion crows (*Corvus corone*), 3% (2/67) thrushes (*Turdus philomelos*), 3% (2/67) Eurasian blue tits (*Cyanistes caeruleus*) and 1% (1/67) of each common chaffinch (*Fringilla coelebs*), common house martin (*Delichon urbicum*), house sparrow (*Passer domesticus*), common swift (*Apus apus*), mallard (*Anas platyrhynchos*) and mute swan (*Cygnus olor*) (Table 3). The confirmed USUV positive animals were 40% (27/67) female and 54% (36/67) male birds; with 6% (4/67) undetermined sex. The USUV positive specimens were 6% (4/67) juveniles, 25% (17/67) subadults and 69% (46/67) adults.

**Table 1a-c.**
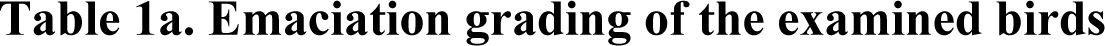
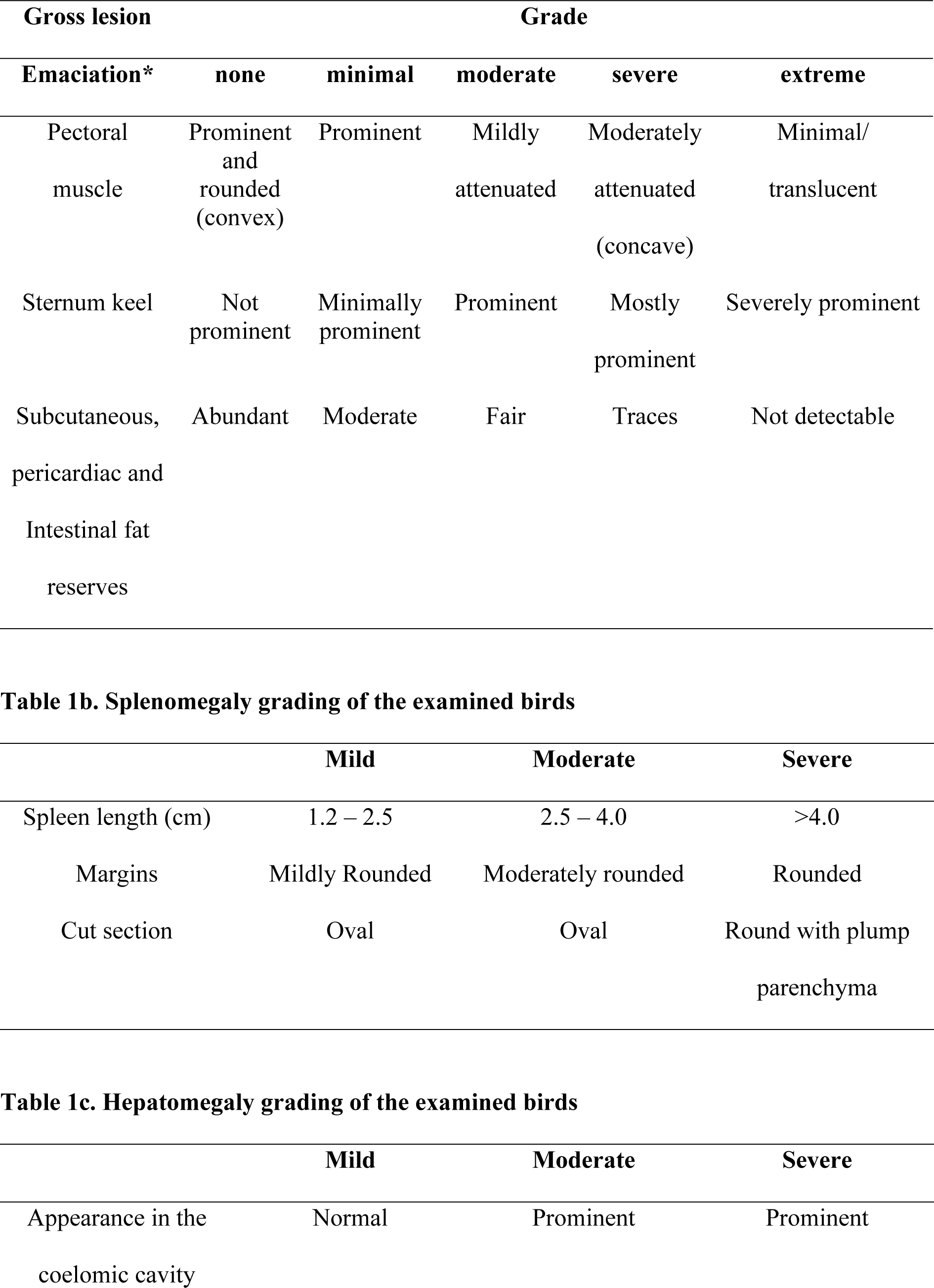
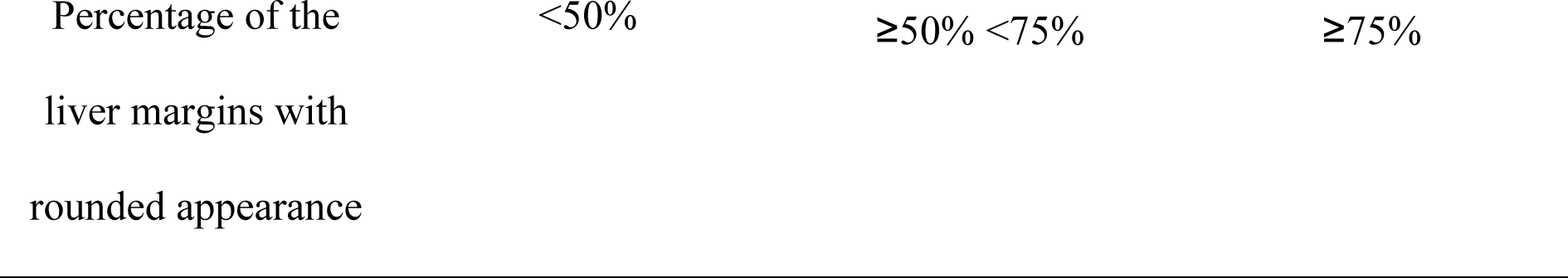
Gross lesions grading of wild birds naturally infected with Usutu virus in Switzerland.

| Gross lesion |  | Grade |  |  |  |
| --- | --- | --- | --- | --- | --- |
| Emaciation* | none | minimal | moderate | severe | extreme |
| Pectoral muscle | Prominent and rounded (convex) | Prominent | Mildly attenuated | Moderately attenuated (concave) | Minimal/translucent |
| Sternum keel | Not prominent | Minimally prominent | Prominent | Mostly prominent | Severely prominent |
| Subcutaneous, pericardiac and Intestinal fat reserves | Abundant | Moderate | Fair | Traces | Not detectable |

|  | Mild | Moderate | Severe |
| --- | --- | --- | --- |
| Spleen length (cm) | 1.2 – 2.5 | 2.5 – 4.0 | >4.0 |
| Margins | Mildly Rounded | Moderately rounded | Rounded |
| Cut section | Oval | Oval | Round with plump parenchyma |

|  | Mild | Moderate | Severe |
| --- | --- | --- | --- |
| Appearance in the coelomic cavity | Normal | Prominent | Prominent |

Percentage of the liver margins with rounded appearance
|  |  |  |
| --- | --- | --- |
| <50% | ≥50% <75% | ≥75% |

**Table 2.**
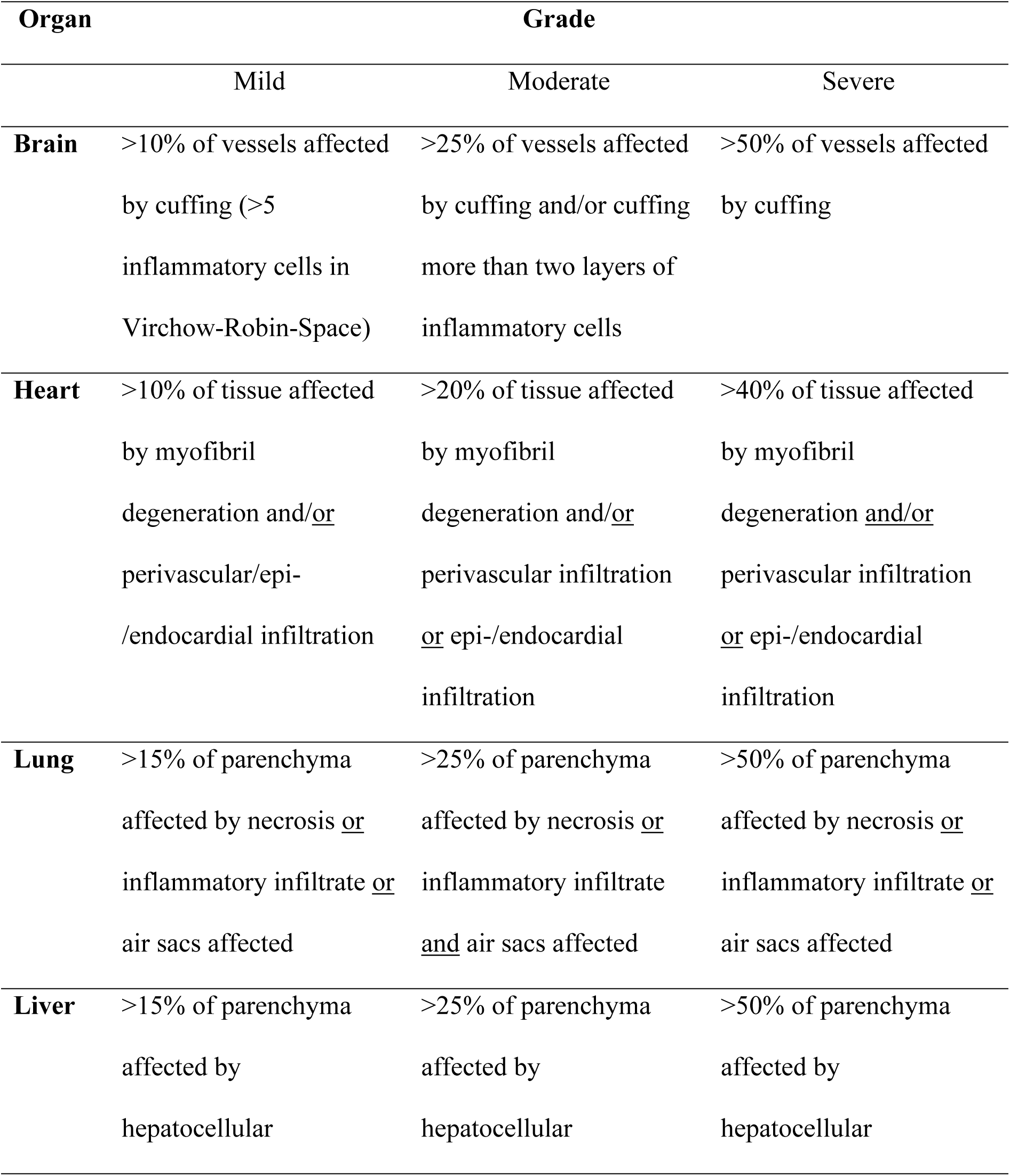
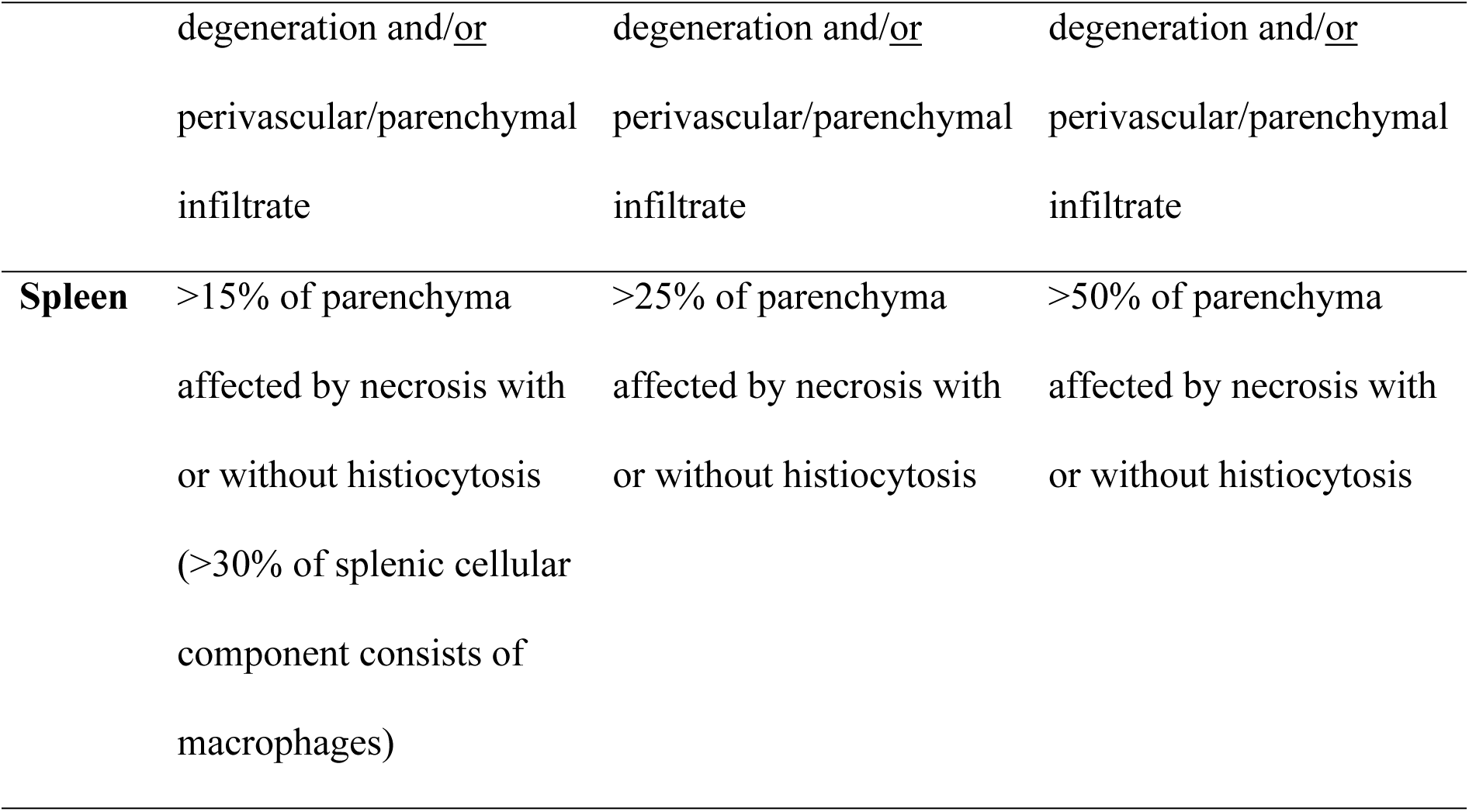
Histological lesions grading of selected organs of wild birds naturally infected with Usutu virus in Switzerland.

| Organ | Grade |  |  |
| --- | --- | --- | --- |
|  | Mild | Moderate | Severe |
| **Brain** | >10% of vessels affected by cuffing (>5 inflammatory cells in Virchow-Robin-Space) | >25% of vessels affected by cuffing and/or cuffing more than two layers of inflammatory cells | >50% of vessels affected by cuffing |
| **Heart** | >10% of tissue affected by myofibril degeneration and/or perivascular/epi-/endocardial infiltration | >20% of tissue affected by myofibril degeneration and/or perivascular infiltration or epi-/endocardial infiltration | >40% of tissue affected by myofibril degeneration and/or perivascular infiltration or epi-/endocardial infiltration |
| **Lung** | >15% of parenchyma affected by necrosis or inflammatory infiltrate or air sacs affected | >25% of parenchyma affected by necrosis or inflammatory infiltrate and air sacs affected | >50% of parenchyma affected by necrosis or inflammatory infiltrate or air sacs affected |
| **Liver** | >15% of parenchyma affected by hepatocellular | >25% of parenchyma affected by hepatocellular | >50% of parenchyma affected by hepatocellular |

|  | degeneration and/or<br>perivascular/parenchymal<br>infiltrate | degeneration and/or<br>perivascular/parenchymal<br>infiltrate | degeneration and/or<br>perivascular/parenchymal<br>infiltrate |
| --- | --- | --- | --- |
| **Spleen** | >15% of parenchyma<br>affected by necrosis with<br>or without histiocytosis<br>(>30% of splenic cellular<br>component consists of<br>macrophages) | >25% of parenchyma<br>affected by necrosis with<br>or without histiocytosis | >50% of parenchyma<br>affected by necrosis with<br>or without histiocytosis |

**Table 3.**
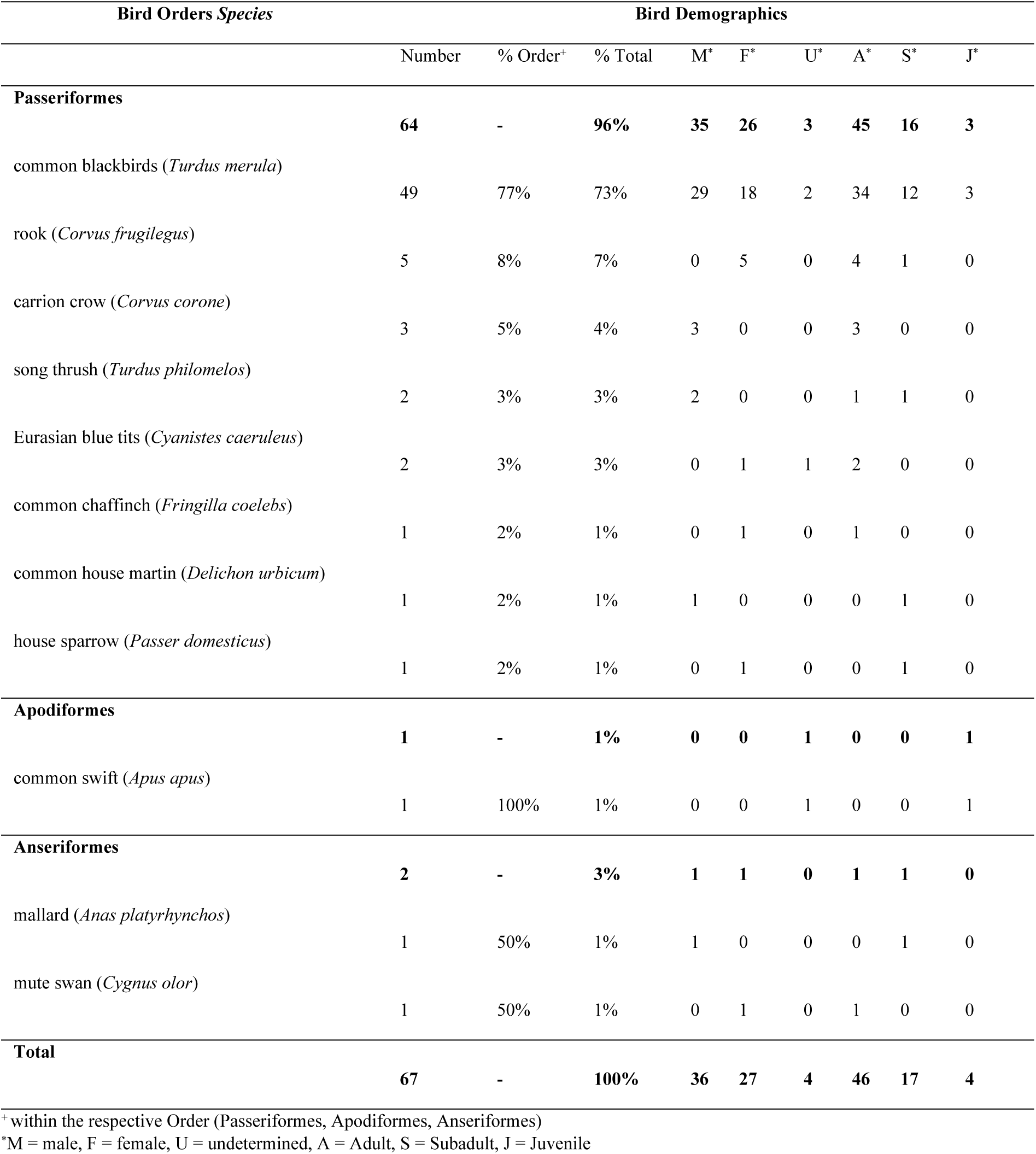
Gender and age distribution of wild bird species naturally infected with Usutu virus in Switzerland as tested by RT-PCR.

| Bird Orders *Species* | Bird Demographics |  |  |  |  |  |  |  |  |
| --- | --- | --- | --- | --- | --- | --- | --- | --- | --- |
|  | Number | % Order<sup>+</sup> | % Total | M* | F* | U* | A* | S* | J* |
| **Passeriformes** | **64** | **-** | **96%** | **35** | **26** | **3** | **45** | **16** | **3** |
| common blackbirds (*Turdus merula*) | 49 | 77% | 73% | 29 | 18 | 2 | 34 | 12 | 3 |
| rook (*Corvus frugilegus*) | 5 | 8% | 7% | 0 | 5 | 0 | 4 | 1 | 0 |
| carriion crow (*Corvus corone*) | 3 | 5% | 4% | 3 | 0 | 0 | 3 | 0 | 0 |
| song thrush (*Turdus philomelos*) | 2 | 3% | 3% | 2 | 0 | 0 | 1 | 1 | 0 |
| Eurasian blue tits (*Cyanistes caeruleus*) | 2 | 3% | 3% | 0 | 1 | 1 | 2 | 0 | 0 |
| common chaffinch (*Fringilla coelebs*) | 1 | 2% | 1% | 0 | 1 | 0 | 1 | 0 | 0 |
| common house martin (*Delichon urbicum*) | 1 | 2% | 1% | 1 | 0 | 0 | 0 | 1 | 0 |
| house sparrow (*Passer domesticus*) | 1 | 2% | 1% | 0 | 1 | 0 | 0 | 1 | 0 |
| **Apodiformes** | **1** | **-** | **1%** | **0** | **0** | **1** | **0** | **0** | **1** |
| common swift (*Apus apus*) | 1 | 100% | 1% | 0 | 0 | 1 | 0 | 0 | 1 |
| **Anseriformes** | **2** | **-** | **3%** | **1** | **1** | **0** | **1** | **1** | **0** |
| mallard (*Anas platyrhynchos*) | 1 | 50% | 1% | 1 | 0 | 0 | 0 | 1 | 0 |
| mute swan (*Cygnus olor*) | 1 | 50% | 1% | 0 | 1 | 0 | 1 | 0 | 0 |
| **Total** | **67** | **-** | **100%** | **36** | **27** | **4** | **46** | **17** | **4** |

<sup>+</sup> within the respective Order (Passeriformes, Apodiformes, Anseriformes)
\*M = male, F = female, U = undetermined, A = Adult, S = Subadult, J = Juvenile

### Pathology

#### Gross findings

Mild to severe loss of condition and mild to severe splenomegaly were the most common gross lesions, which were observed in 84% (56/67) and 75% (50/67) of the USUV positive birds by RT-PCR, respectively. Other findings compatible with Usutu virus disease (UVD) were mild to severe hepatomegaly in 12% (8/67) and multiple-organ congestion in 10% (7/67) of the RT-PCR positive animals. Additional, accidental findings included signs of trauma (hemorrhages, lacerations, perforations, ruptures, fractures) in 21% (14/67) of the examined birds. Intestinal endoparasites (nematodes and cestodes not further characterized) and/or trematodes within the air sacs (Family *Cyclocoelidae*) were observed in 51% (34/67) of the animals. Feather abnormalities were observed in 33% of the birds (22/67), including persistent feather sheaths (19%, 13/67), loss of feathers at the head, wings or tail (12%, 8/67) and short feathers (4%, 3/67). Ninety-one percent (20/22) of birds with feather abnormalities were common blackbirds.

#### Histopathology

Histopathology could be evaluated on 34 of the 67 confirmed USUV positive animals and a homogeneous set of organs (brain, heart, lungs, liver, kidney, spleen) could be examined in no less than 26 out of 34 animals (76%).

Sixty-five percent (22/34) of the birds showed histological lesions in the brain (Figure 1), with perivascular cuffing (n=14), cuffing and gliosis (n=1), meningitis only (n=2) and cuffing and meningitis (n=2), and gliosis only (n=3). The changes were generally mild, with one exception consistent with severe encephalitis in one blackbird.

**Fig. 1:**
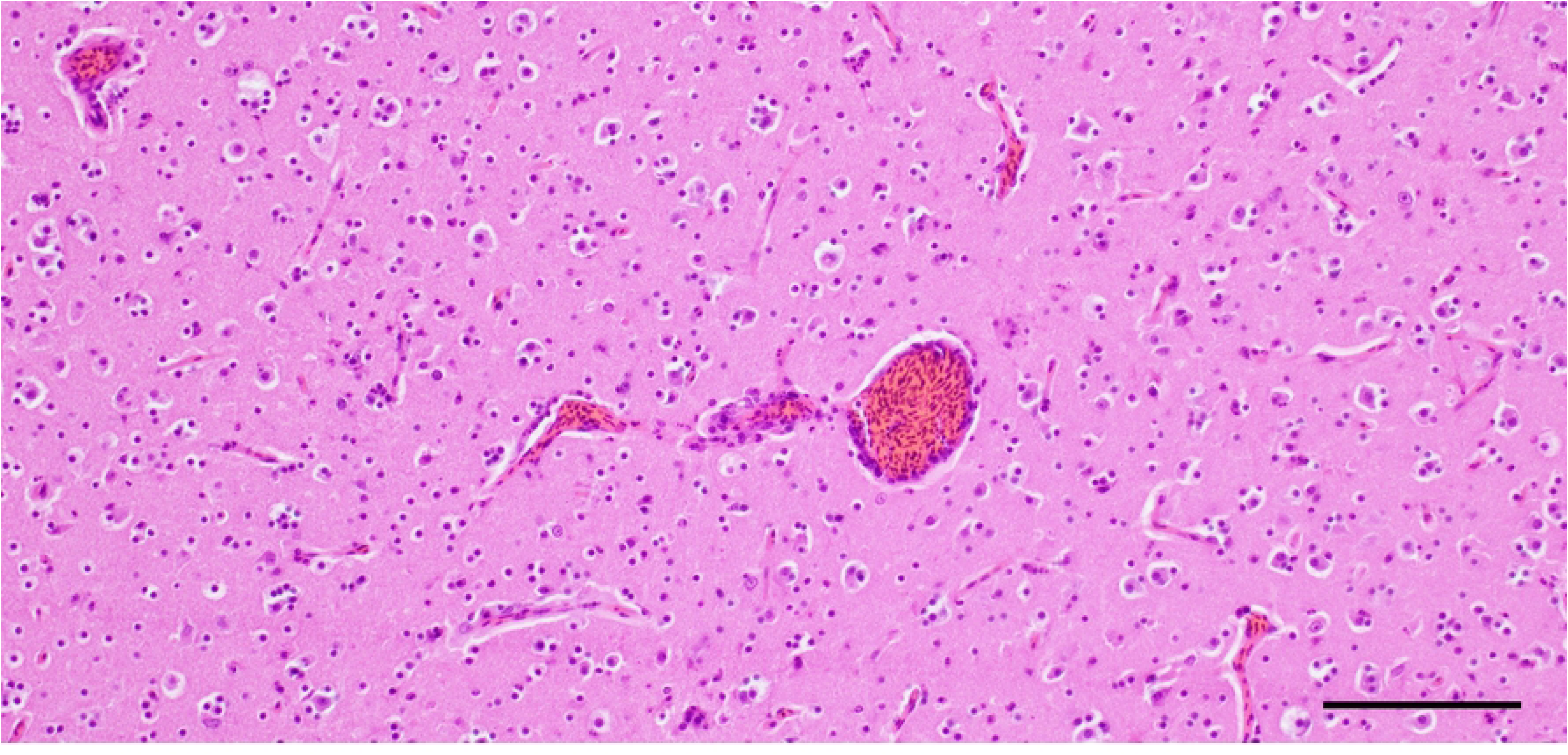
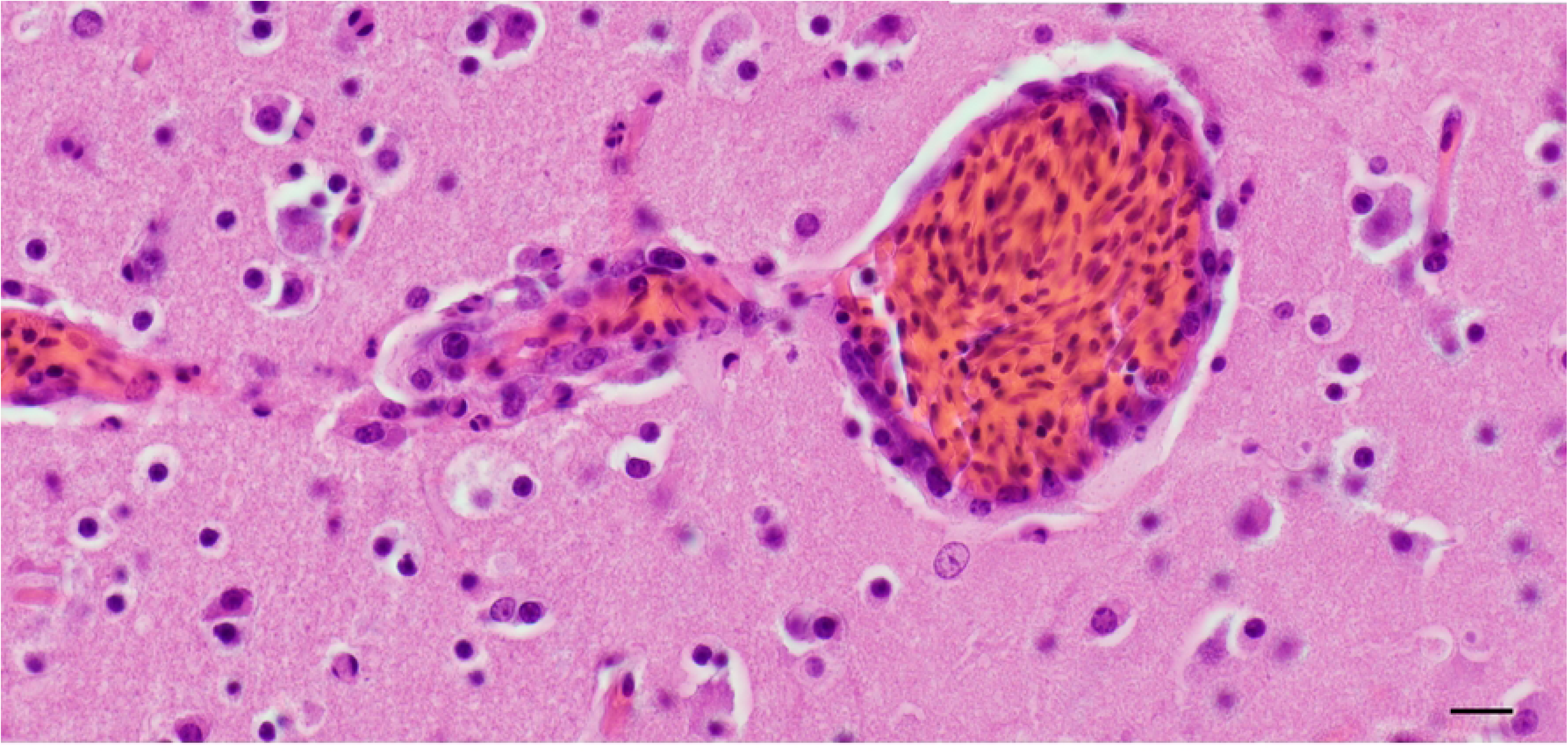
Representative photomicrograph of a brain section of a blackbird (*Turdus merula*) with Usutu virus infection and associated inflammation (encephalitis). (Left) The Virchow-Robin space is expanded by an inflammatory infiltrate (perivascular cuffing) along with reactive endothelium (endothelial hypertrophy). Bar 100 um. H&E. (Right) The infiltrate is composed of mainly macrophages and fewer lymphocytes. Bar 20 um. H&E.

Fifty-six per cent (19/34) of examined animals presented cardiac lesions (Figure 2), ranging from minimal to moderate lympho-histiocytic myocarditis (n=13), epicarditis with myofibril degeneration and loss (n=2), combined lympho-histiocytic myocarditis and lympho-plasmacytic endocarditis (n=3), and pancarditis (n=1). The inflammatory process was mostly, but not exclusively, centered on perivascular spaces.

**Fig. 2:**
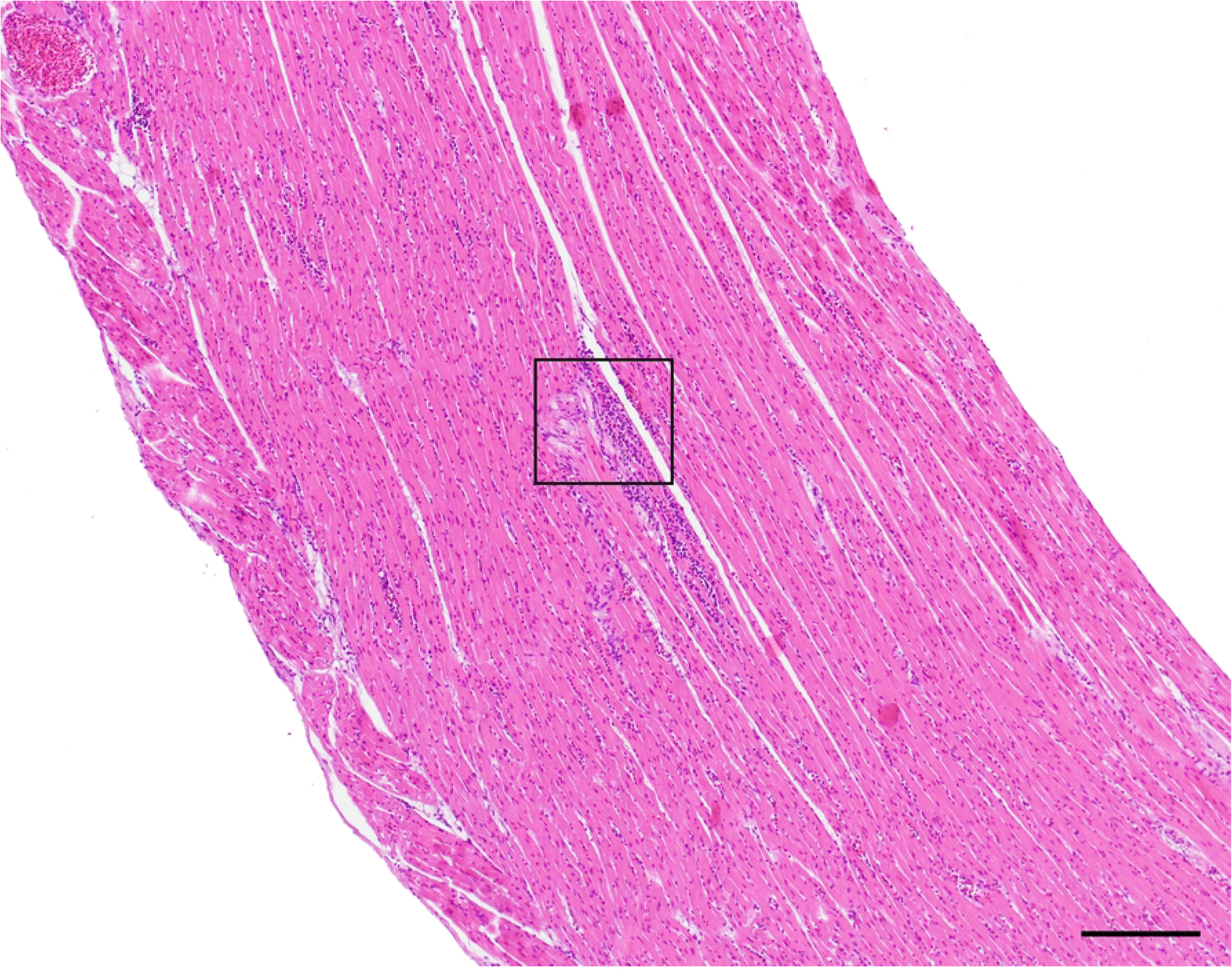
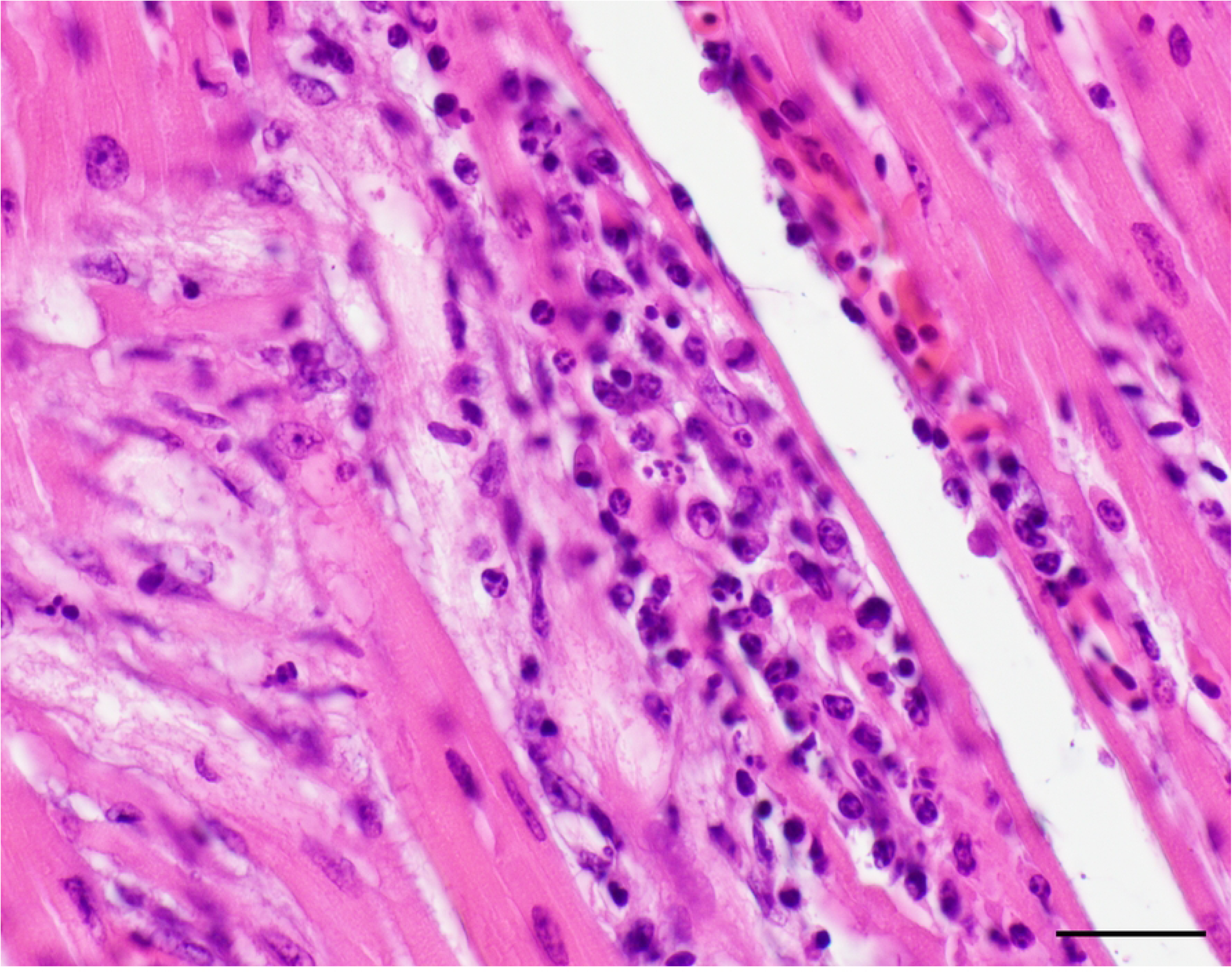
Representative photomicrograph of a heart section of a blackbird (*Turdus merula*) with Usutu virus infection and associated inflammation (myocarditis). (Left) A mixed inflammatory infiltrate expands the interstitium, bordering the interfibril clear spaces (edema). Bar 500 um. H&E. (Right) The infiltrate is composed mainly of macrophages, fewer heterophils and plasma cells with admixed degenerate myocardial fibers. Bar 50 um. H&E.

Tissue changes were present in 62% (21/34) of the examined liver sections (Figure 3). Birds showed mild to moderate lympho-histiocytic infiltrates (n=19) in the liver parenchyma, frequently associated with multiple necrotic foci/hepatocellular loss (n=13) and two with multifocal hepatic necrosis only (Figure 3).

**Fig. 3:**
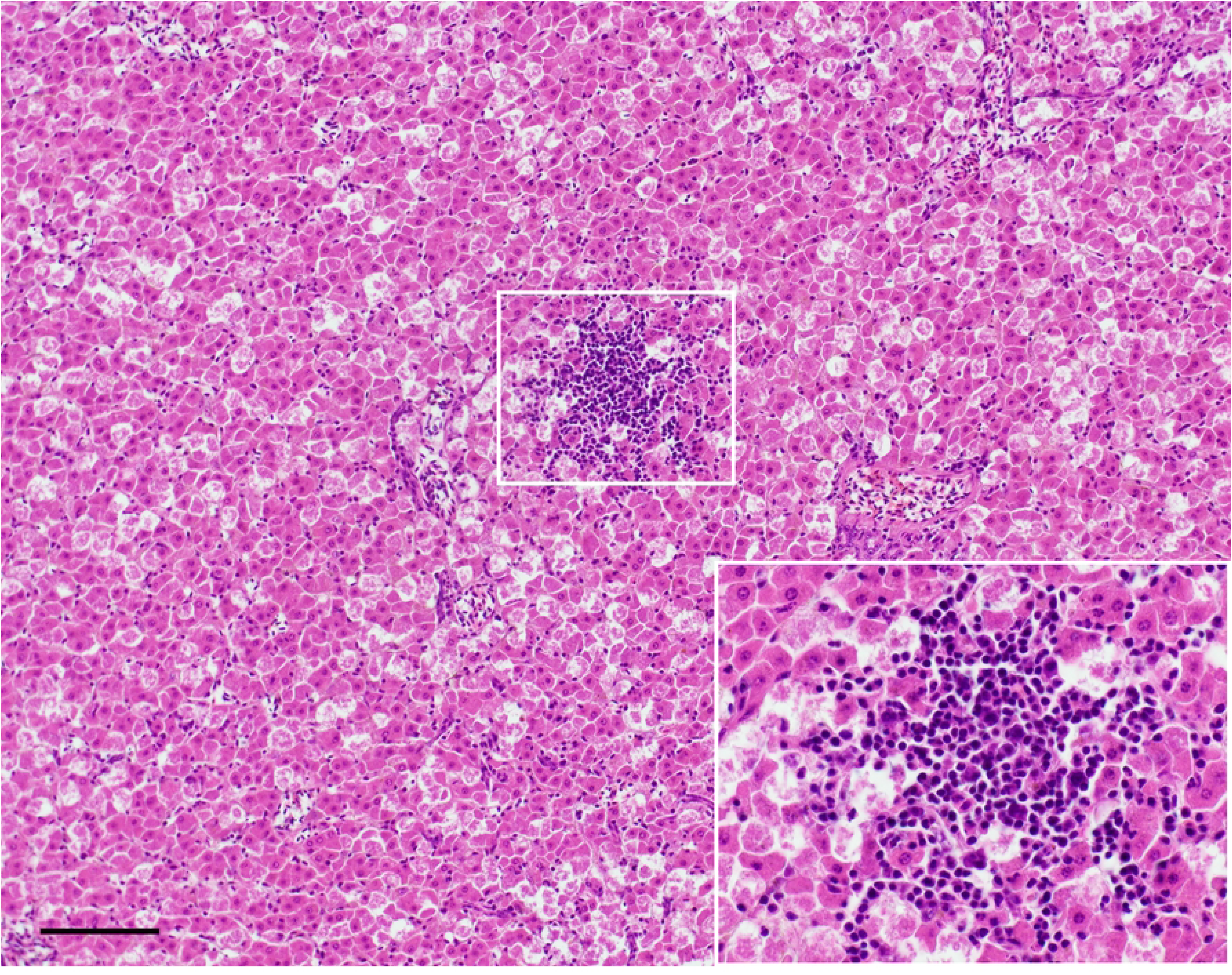
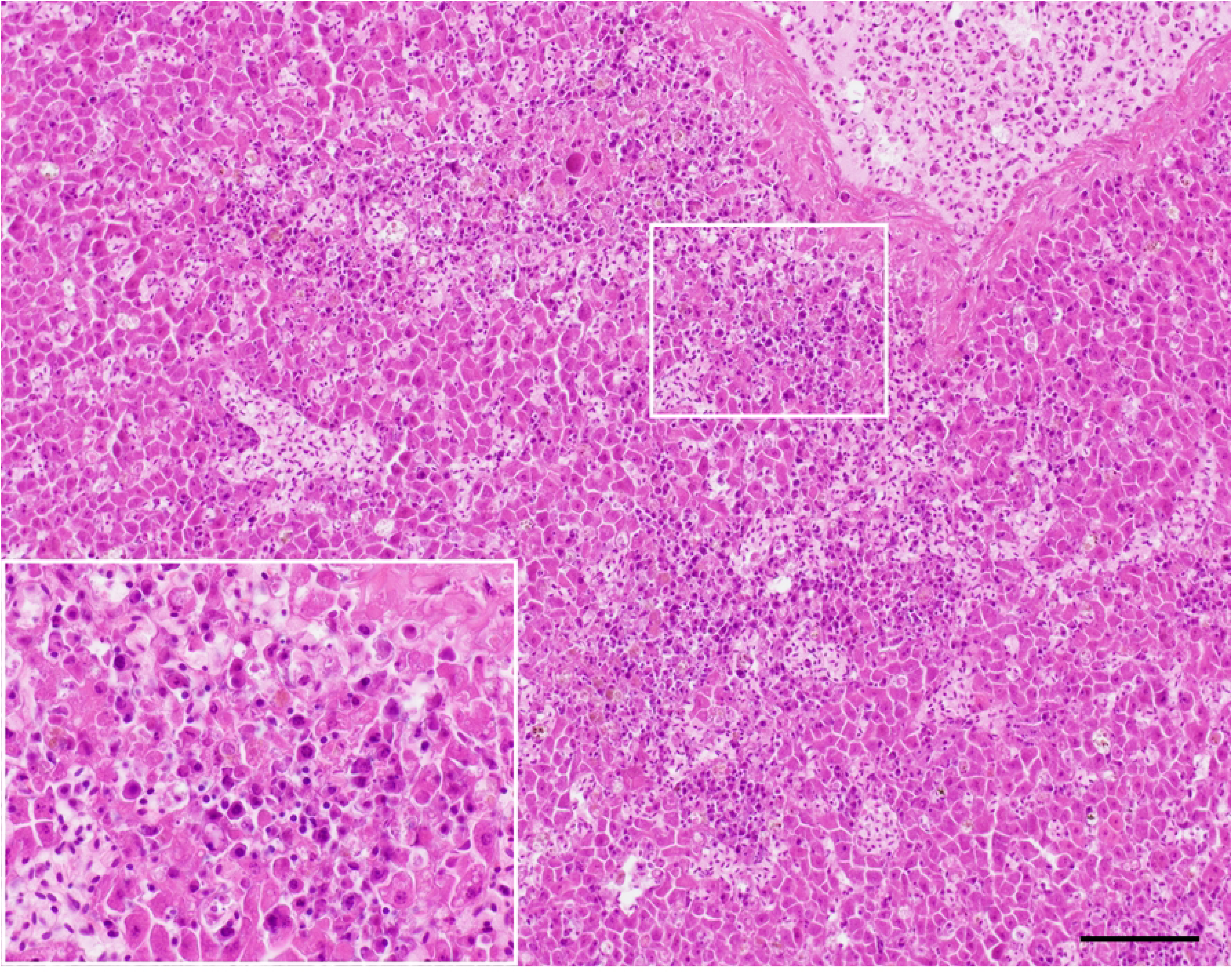
Representative photomicrograph of liver sections in a blackbird (*Turdus merula*) with Usutu virus infection and associated inflammation (hepatitis). (Left) An inflammatory infiltrate is separating and replacing hepatocytes. The infiltrate is mainly composed of plasma cells and lymphocytes. The inset is a magnification of the marked area. Bar 100 um. H&E. (Right) Multifocal to coalescing areas display loss of hepatocytes. Hepatocytes are either swollen and pale (degenerating) or shrunken and hypereosinophilic (necrotic). The inset is a magnification of the marked area. Bar 100 um. H&E.

Eighty-six percent of the examined spleens (25/29) showed three characteristic arrangements of the infiltration of histiocytes. The histiocytic infiltrate could be seen both as organized in (1) discrete clusters (n=11) and as (2) dispersed (n=25) within the spleen parenchyma. Hereby discrete clusters could always be observed in the same tissue section along with dispersed histiocytes. Loosely grouped macrophages (histiocytic aggregates) were accompanied by large numbers of plasma cells that floated the splenic parenchyma (Figure 4A). Later on, discrete clusters of macrophages became more apparent (Figure 4B). Finally, when (3) dispersed histiocytes were degenerated and necrotic we observed them together with severe engorgement of the spleen by erythrocytes (observed as prominent splenomegaly) and progressive depletion of the white pulp (Figure 4C). Additionally, parenchymal necrosis was relatively commonly observed (n=21).

**Fig. 4A:**
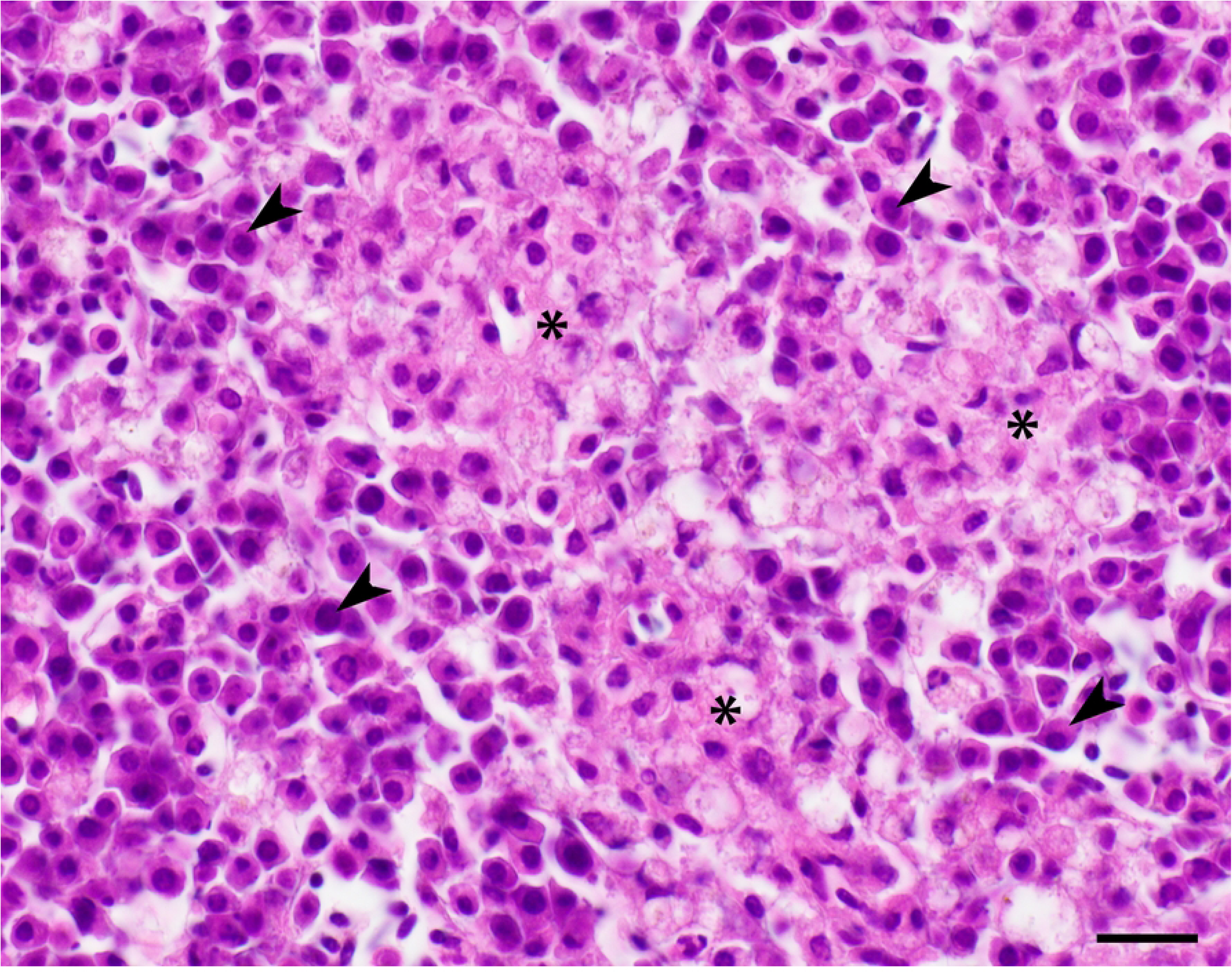
Representative photomicrograph of spleen sections in a blackbird (*Turdus merula*) with Usutu virus infection and associated presumptive **early splenic changes**. (Left) Dispersed macrophages are beginning to form clusters visible as pale areas (asterisks). Bar 200 um. H&E. Large number of plasma cells are floating the parenchyma (arrowheads). Macrophages begin to cluster (asterisks). Bar 20 um. H&E.

**Fig. 4B:**
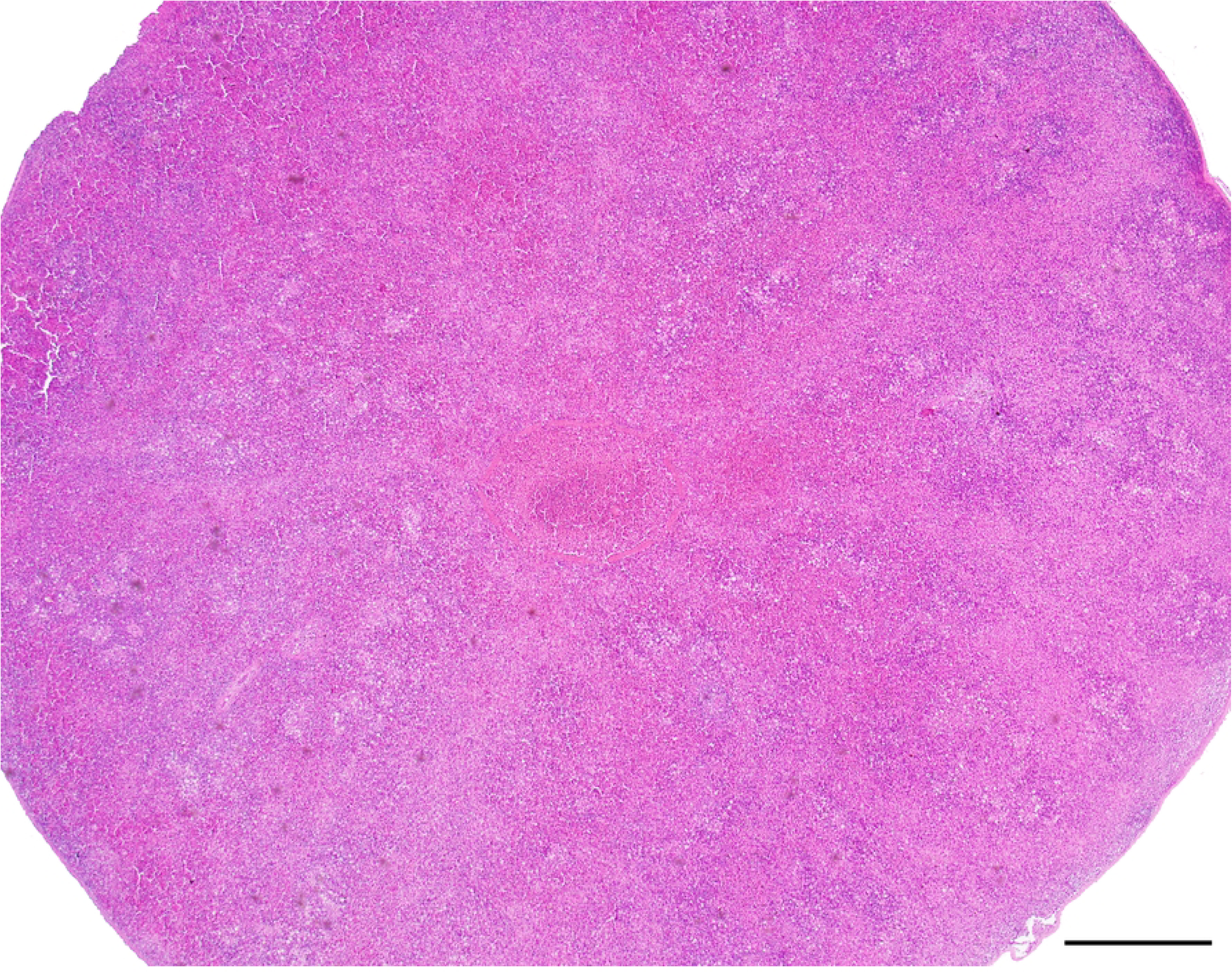
Representative photomicrograph of spleen sections in a blackbird (*Turdus merula*) with Usutu virus infection and associated presumptive **intermediate splenic changes**. (Left) The spleen is markedly swollen (round) and macrophages clusters are clearly visible as pale areas. Bar 500 um. H&E. Macrophages are clusters together (arrowheads). Bar 50 um. H&E.

**Fig. 4C:**
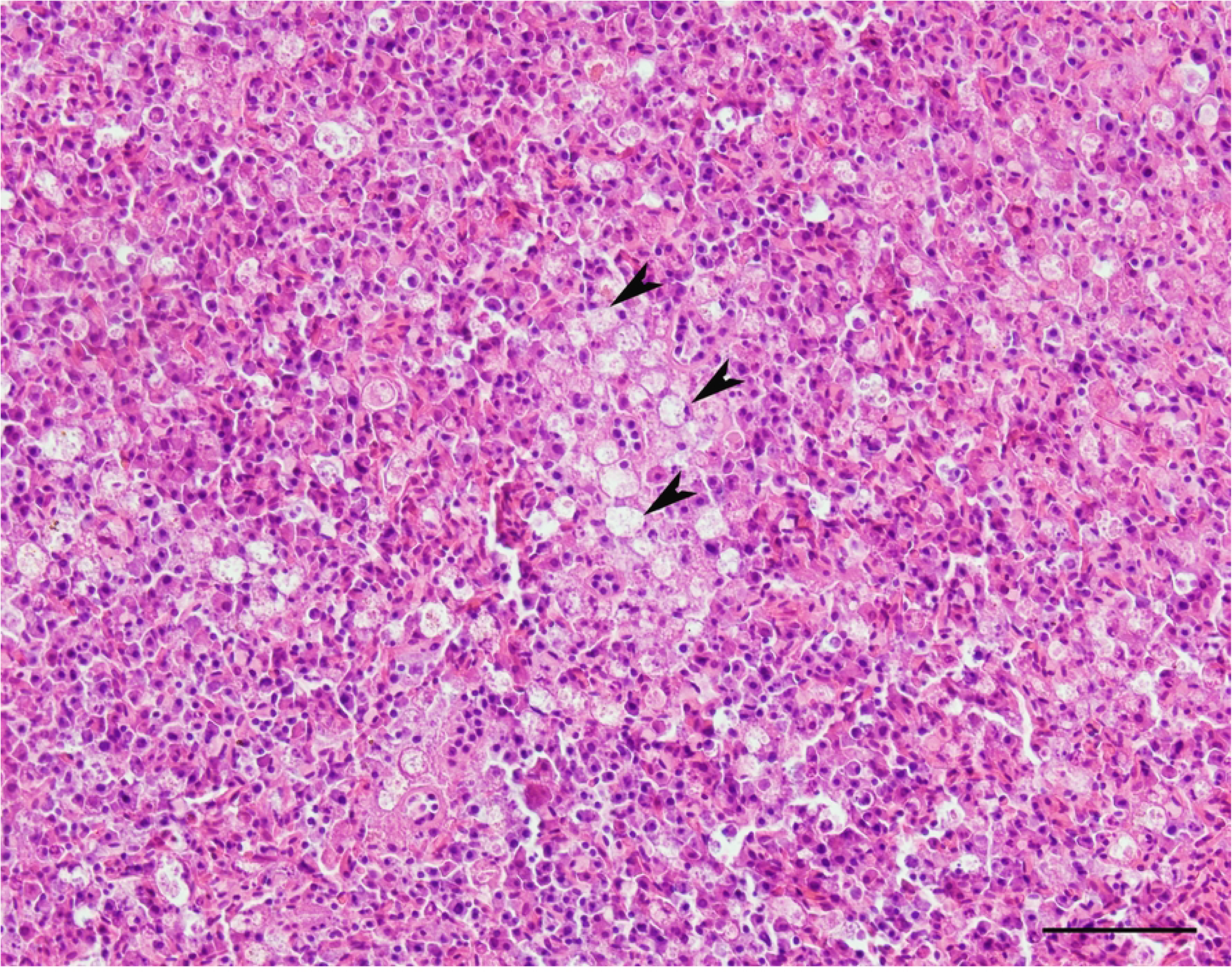
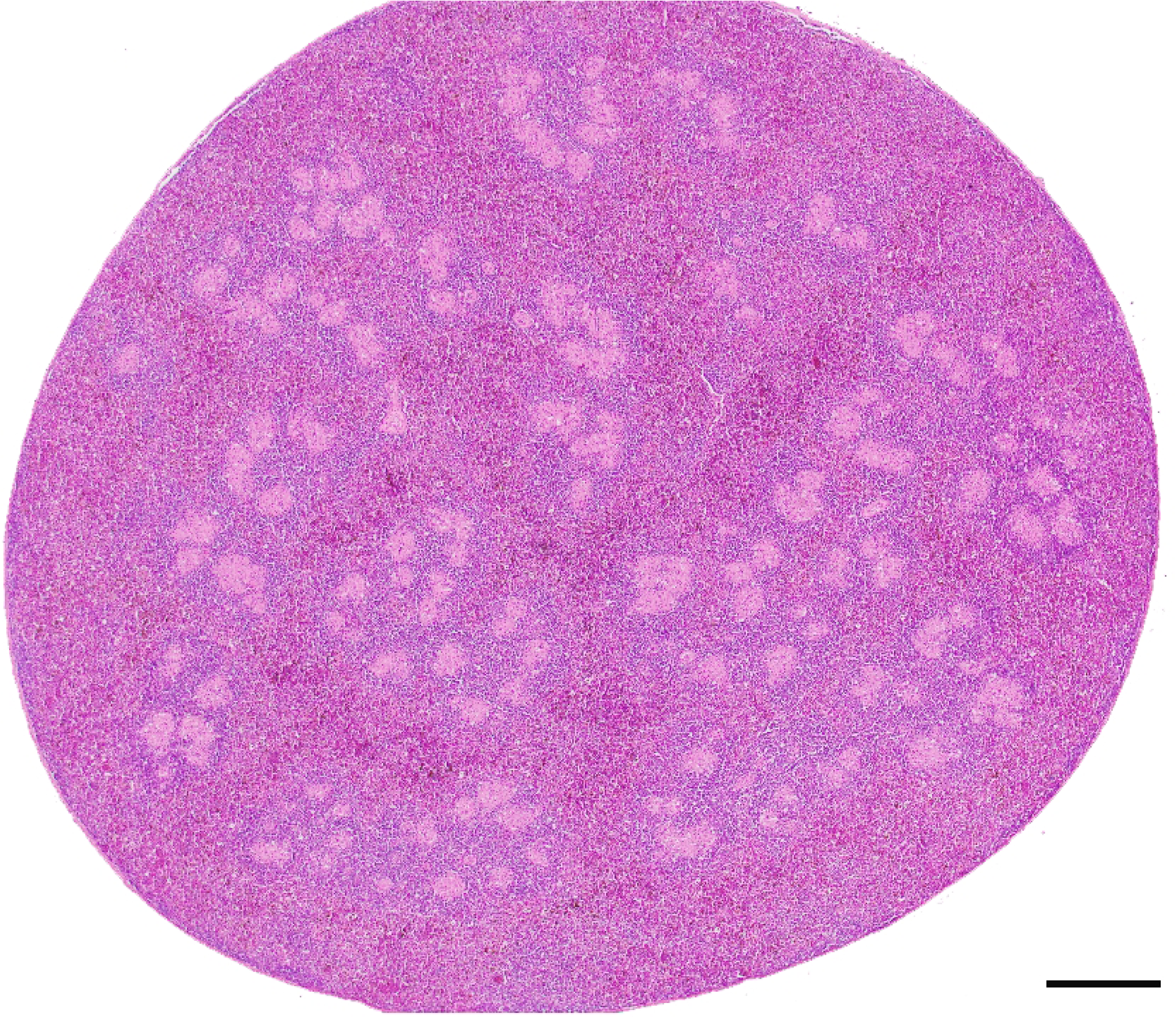
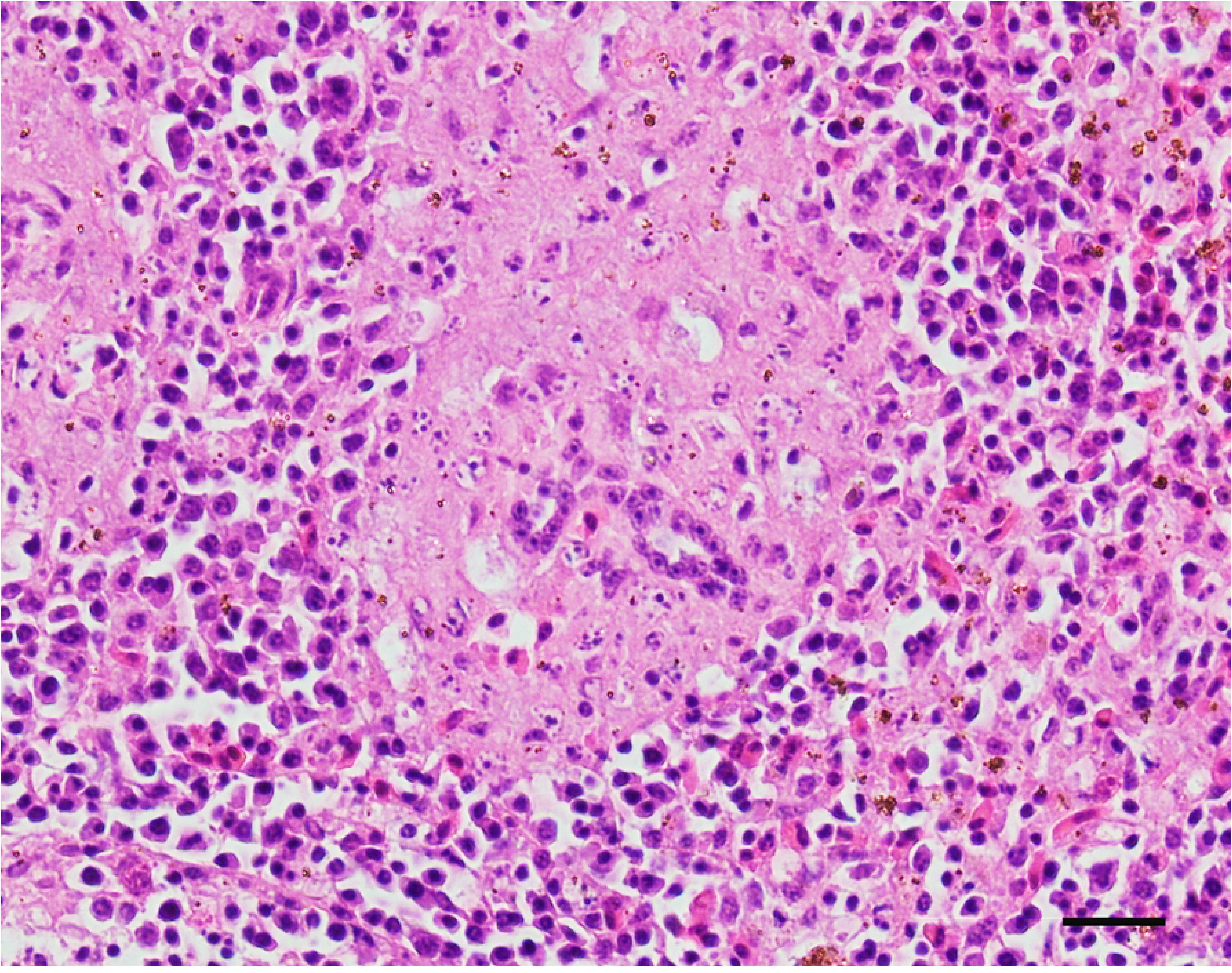
Representative photomicrograph of spleen sections in a blackbird (*Turdus merula*) with Usutu virus infection and associated presumptive **late splenic changes**. (Left) The spleen is visibly enlarged (round). There is marked accumulation of erythrocytes (congestion) and a lack of lymphoid follicles (lymphoid depletion). Bar 500 um. H&E. (Right) Lymphocytes depleted ellipsoids and pennicillar arterioles are prominent and show the naked surrounding collagen. Bar 20. H&E.

Additional lesions included respiratory tract changes in 68% (23/34) of the animals consistent with mild to moderate either interstitial or granulomatous pneumonia (n=10), lympho-histiocytic to plasmacellular aerosacculitis (n=3), a combination of both (n=7) or bronchitis (n=3). In 26% (6/23) of the lung lesions, the inflammatory process was associated with an obvious presence of infectious and/or infestive agent, i.e. fungi (n=3) or nematode larvae (n=3). Twenty per cent (7/35) of the birds showed mild to moderate pneumoconiosis. Four birds (13%; 4/32) had mild tubular degeneration (n=1) and lympho-histiocytic to heterophilic interstitial nephritis (n=3).

### RT-PCR and phylogenetic analysis

Forty-nine of the 67 RT-PCR positive birds yielded a readable nucleotide sequence corresponding to part of the NS5 region reported to be a reliable phylogenetic proxy for the entire viral genome (Cadar et al. 2017a; Engel et al. 2016). These samples, which are those unambiguously proven USUVS positive were those chosen for the investigation presented here. Of these, the majority (96%; 47/49) clustered together with the USUV lineage Europe 3 (Figure 5A), with 53% (26/49) sharing 98.58 to 100% identity with USUV-BE-Angleur strain (KY263623.1), 29% (14/49) 100% matching the USUV strain USUV-BE-Flemalle (KY263624.1) and 14% (7/49) sharing 99.53 to 100% identity with USUV strain USUV-BO-8 (HM138716.1) (Table 4). Nine Swiss USUV strains showed silent single nucleotide substitutions (SNP) of the amplified partial NS5 region when compared to the reference strain (1 SNP in 7 Angleur-like strains, 1 SNP in 1 Bonn-like strain and 2 SNPs in 1 Angleur-like strain). One Angleur-like strain showed a single nucleotide deletion associated with a frameshift mutation (S1 Table).

**Fig. 5A:** Phylogenetic tree of a representative selection of Usutu virus strains compared to the closely related West Nile Virus. In boxes are the strains detected in this study. Highlighted are different lineages. The majority of strains in this study appear to cluster within the lineage Europe 3. However, two samples are part of the lineage Africa 3. **5B**: Bar chart of identified strains of Usutu virus in Switzerland in wild birds. No strain was successfully sequenced in 2020. **5C**: Map of Switzerland with distribution of different Usutu virus strains. Visible are Flamelle in turquoise, Bonn in green, BO-8 in magenta, Angleur in yellow (non-sequenced samples are shown as dark grey circles).

**Table 4.**
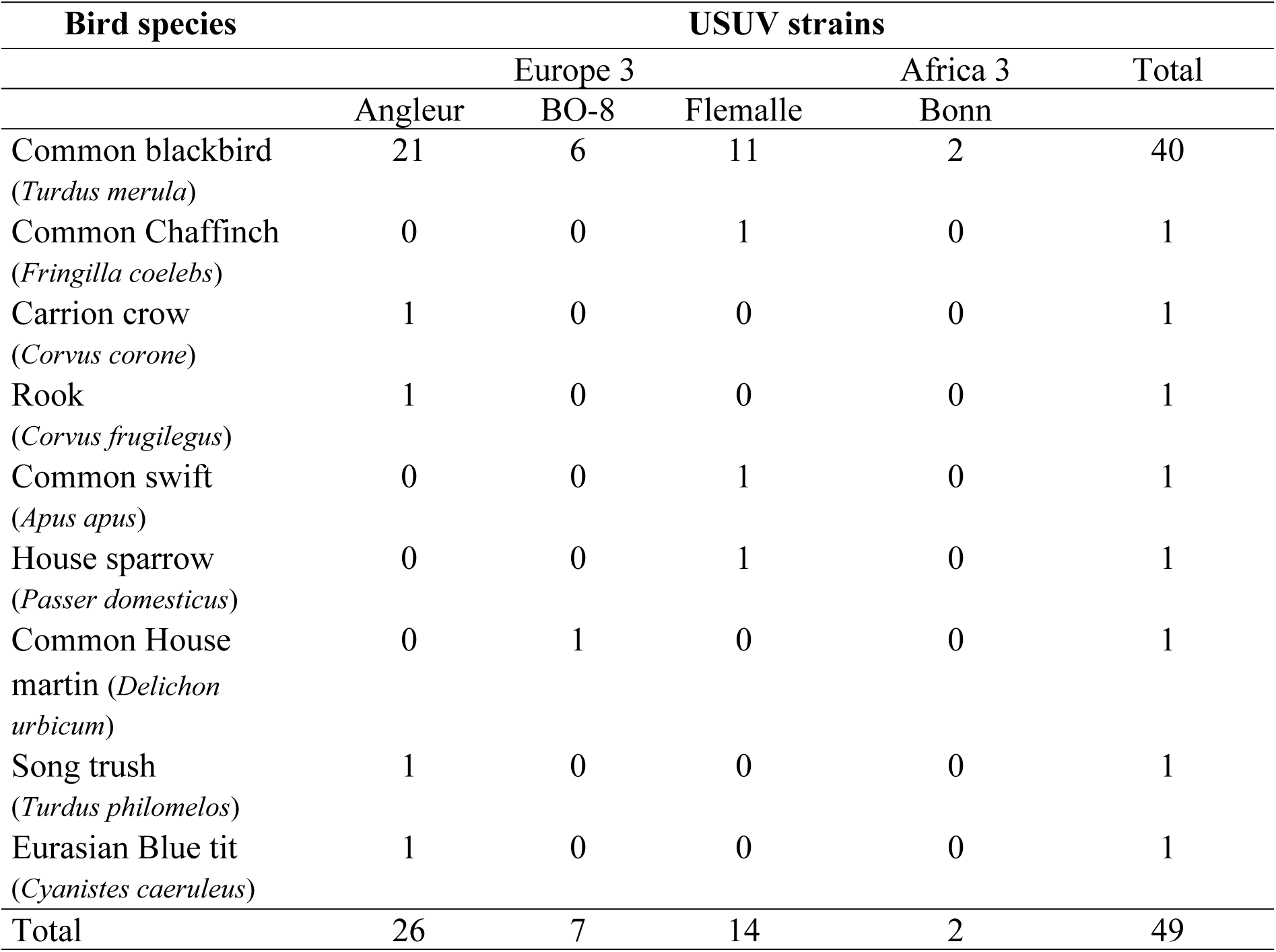
Wild bird species distribution of lineages of USUV infection.

The remaining two USUV lineage (4%) were Africa 3 strains were found in two blackbirds from the Southwestern Alps and were 100% identical to the USUV strain Bonn (KM659877.1). Alignment is available as supplementary material.

The bulk of the obtained sequences (76%; 37/49) were sampled in 2018, reflecting an obvious infection spike within the time span considered (2015-2020). Accordingly, the remaining samples were relatively similarly distributed prior and after 2018, with 4% (2/49) in 2015, 2% (1/49) in 2016, 12% (6/49) in 2017, and 6% (3/49) in 2019 (Figure 5B). Angleur-like strain was the first one to be detected in 2015 in two black birds and in the following year in a rook. In 2017 two new variants, specifically Flemalle- and BO-8-like strains, were detected in a few blackbirds (n=6). The year 2018 was the only time when USUV strains of the lineage Africa 3 were detected in black birds, whereas no lineage other than Europe 3 was detected during the remaining years. Another outstanding feature of the 2018 epidemic was the increased number of bird species infected with USUV. Beside of the overrepresented blackbird, all three strains of lineage Europe 3 (known to be present in Switzerland to date) were isolated in house sparrows, common chaffinch, song thrush, Eurasian blue tit, carrion crow and common house martin (Table 4).

### Phylogeography

We received submissions for potential USUV cases from 17 of the 26 Swiss cantons and from the Principality of Liechtenstein, with most of the cases from North and Central Switzerland and fewer submissions from South and East Switzerland. Accordingly, the RT-PCR confirmed positive animals were mainly distributed within the Plateau (91.0%, 61/67), corresponding to North-Central Switzerland and then equally within the Jura (4.5%, 3/67) (North-Western) and within the Alps (4.5%, 3/67) (South-Eastern) (Figure 5C).

Birds infected with USUV Europe lineages (Angleur-, Flemalle- and BO-8-like strains), were above all from the Plateau (North Central to West Central) and to a lesser extent from the Jura (North West). No outstanding variation in the distribution of Flemalle- and Angleur-like strains were observed (mostly Plateau and lesser in the Jura), however there was an apparent clustering of BO-8-like strains between the two metropolitan areas of Zurich and Luzern (North central, Plateau). USUV Africa 3 strains were detected exclusively from southern Switzerland (Canton Valais). Accordingly, the phylogeographic analysis of the USUV strains circulating in Switzerland reveals a predominant distribution of diverse Europe lineages (mainly Europe lineage 3) across the country, at the exception of the Africa 3 lineage, which is the only lineage that could be found in the Alps, in the Southwestern portion of the country.

### Environmental Conditions and Vectors

According to our observations and results, 2018 was the year when the large majority of the confirmed positive USUV cases occurred. Accordingly, special attention was paid to possible outstanding environmental conditions, which might have occurred in 2018 and eventually primed the surge of USUV cases. Among others, meteorological conditions, known to be determinant in influencing the dynamic of the USUV vector populations (mainly *Culex* spp.) as already studied for other arboviruses [48,49], were carefully considered. According to the Swiss meteorological service (Federal Office of Meteorology and Climatology MeteoSwiss) the years 2014 to 2018 were the warmest five years ever recorded [50]. In particular, 2018 was the fourth warmest year (0.3°C above the 1981– 2010) since measurements began in 1864 and a high annual average temperature was recorded on almost all continents. The same institution pointed out that the year 2018 was, with +1.5°C, the warmest year and the third warmest Summer in Switzerland since measurements began in 1864 [50,51]. The national average for April was 3.9°C above the average. Summer 2018 was the third year in a row with an above-average temperature with a deviation of 2.0°C. The hot summer was followed by the third warmest autumn since the start of measurements in 1864. The precipitation and humidity in spring to autumn were consistently lower (drier weather) than the average. In the eight months from April to November only 59% of the average precipitation occurred.

Although no comprehensive information of direct evidence of an abnormal increase of the *Cx. pipiens* population, the USUV vector, in the summer of 2018, was available, indirect evidence was provided by the epidemiological data concerning another flavivirus, West Nile Virus (WNV). More specifically, *Cx. pipiens* serves as a vector not only of USUV but also of WNV. Strikingly, the World Health Organization (WHO) reported that WNV cases underwent a sharp increase in 2018 if compared with the previous 4 years. The WHO underlines that “*This is largely due to the early start of the 2018 transmission season in the WHO European Region, which normally lasts from July to October. This year’s season <u>(2018, note of the authors</u>) has been characterized by high temperatures and extended rainy spells, followed by dry weather. Such weather conditions have been conducive to mosquito breeding and propagation*”. [52] Accordingly, the Swiss and panEuropean surge of USUV in 2018, might have had a similar background, since the season was hot during late spring and early summer, which favored the increase of mosquitoes.

## Discussion

This investigation aimed to explore the features of Usutu virus-associated disease in free ranging birds in Switzerland from 2015 to 2020, contributing to highlight those factors, which might have been relevant for the occurrence of the major outbreak of 2018. Forty-nine birds were confirmed conclusively USUV positive by RT-PCR and sequencing. USUV is known to affect mainly birds of the order Passeriformes, with common blackbirds as the most affected species. Consistently, 73% (49/67) of the USUV positive birds were blackbirds. Additionally, our investigation is the first to report confirmed USUV infection with associated lesions in new species (*Corvus frugilegus*). Furthermore, we provided the first USUV infection in a mute swan (*Cygnus olor*), a species for which only exposure to USUV by the presence of antibodies, was reported up to now (Meister et al. 2008). Unfortunately, lineage could not be detected in the swan and histopathology was not performed. Moreover, although USUV has just recently been detected and described in common house martin (*Delichon urbica*), common swifts (*Apus apus*), common chaffinches (*Fringilla coelebs*) and mallard ducks (*Anas platyrhynchos*), associated histological findings were not described [24,53]. Differently, in our study the common swift and the common chaffinch presented classic USUV associated lesions, including spleen and kidney tubular necrosis (common swift) together with gliosis and with mild myocarditis (common chaffinch). Interestingly, the USUV positive mallard showed no specific pathological findings in the major organs examined (brain, heart, lung, liver, kidney, and spleen). Similar to many of the submitted birds, the mallard was found dead, and it is not clear if the diseases progressed hyperacutely in this bird, preventing the development of detectable lesions, or if they were too mild to be differentiated from the commonly observed background autolysis. Overall, our findings further expand the known USUV host spectrum, suggesting that several additional species, besides those traditionally known, may be susceptible to the viral infection and eventually to the development of the disease.

According to the examined birds, we observed slightly more positive males than females (54% male vs. 40% female birds). However, these data need to be interpreted cautiously because a likely sampling bias cannot be ruled out secondary to the more readily detectable male blackbird (by far the most represented bird species) in the field because of their darker feather coat and bright yellow beak. Interestingly, we observed that the proportion of young individuals examined (21%) was not associated with a proportional number of positive individuals (8%), suggesting that infection may occur more commonly in adult birds (data not shown). Further studies are needed to confirm these initial observations.

In line with previous reports [10,11,54] we observed that occurrence of UVD in Switzerland showed unspecific gross pathology signs, consistent with an often severe, splenomegaly together with poor body condition, (based on detectable fat deposits and pectoral musculature development), most frequently observed in blackbirds. A gross finding with questionable relation to USUV infection, but frequently observed in submitted black birds, were feather abnormalities consistent with persistent feather sheaths, featherless heads, and loss of feathers at the wings or tail, which we observed from August till November. Other studies, e.g. from the Netherlands [17] described the same phenomenon. Physiological molt for blackbirds is described from June till October but happens mostly in August. Accordingly, we interpreted these findings as most likely a part of a physiological process but unusually extended in time. It is tempting to consider an alteration of the normal molting process eventually associated with the USUV infection, with the virus presumptively damaging directly or indirectly the feather follicles and their cycling.

Histological lesions which have been previously described in UVD included mainly myocarditis, encephalitis, splenitis, hepatitis, as well as splenic and hepatic and, occasionally, renal necrosis [10,11,35,55,56]. More specifically, a study from Germany reported that spleen and liver represent potential early sides of viral infection as shown with immunochemistry for double-stranded ribonucleic acid [57]. In this study histological lesions associated with confirmed USUV infection were predominantly observed in the heart, brain, liver and spleen, consistently with what is described in the literature [5,8,33,35].

Of particular interest was the close examination of the microscopic changes detected in the spleen, the organ, which was more consistently grossly affected in the infected birds, especially in blackbirds. Interestingly, three distinct patterns including (1) compact multifocal, dense clusters of histiocytes with or without necrosis, appeared to expand to form (2) larger, loose aggregates more homogenously distributed in the whole spleen parenchyma, finally obscured by a (3) massive engorgement of erythrocytes in a severely lymphoid-depleted spleen. To our understanding these pictures could mirror different stages of the pathological process within the spleen during an USUV infection (early, intermediate and late splenic changes, Fig. 4A-C). Histiocytes are notoriously pivotal elements in the antiviral response [58] and commonly pathogen-hijacked immune cells, which often our initial observations are highly suggestive of a crucial role of histiocytes in the pathogenesis of USUV associated disease. In line with this are several studies detecting viral antigen in macrophages in different organ systems via immunohistochemical labeling [54]. The question remains open if this is part of the innate immune response and viral phagocytosis or if macrophages are specifically targeted by the virus and used for replication as seen for WNV [59–61]. Presumptively, an initial invasion of USUV into the splenic parenchyma might be associated with the formation of clusters of histiocytes or activated macrophages within normal lymphoid cells and splenic architecture. Later on, those clusters would lose integrity and macrophages disperse into the lymphoid tissue, partially replacing it. The progressive necrosis and loss of the macrophages, possibly, because of the direct effect of the virus (likely replicating into the macrophage cytoplasm), would deplete the spleen parenchyma of these cells and would be followed by the engorgement of the red blood cells, the most likely main responsible of the end stage splenomegaly observed in affected birds. Whether this process or these stages are classified as histiocytosis, histiocytic splenitis or an early stage of granulomatous splenitis needs to be discussed, as to date this aspect of USD is lacking investigations. Why some infected birds (17/67) did not show splenomegaly, remains to be determined. Possible explanations include among others, individual immune response, an early stage of infection or different USUV strains associated with distinct pathology phenotypes. Curiously, the most well defined and discrete macrophages clusters were observed in relatively small spleens, consistent with the collection of these groups of cells in the early stage of the disease. In our sample collection, almost all Bonn-like strains BO-8- and Flemalle-like strain infected birds were associated with splenomegaly. On the other hand, only three fourth of the birds infected with the Angleur-like strain showed splenomegaly. Our sample size is relatively small to draw definitive conclusions, however it does not appear unrealistic the existence of a possible molecular signature associated with the infecting strains translating into distinct pathology phenotypes, as known for other infectious agents [62]. Important is the high number of lesions we observed in the respiratory system. Pulmonary pathology has only rarely been described in UVD [5,24]. Even excluding lesions where obvious infectious agents were present, the high percentage of birds with aerosacculitis and/or pneumonia (77%; n=23/30) might suggest that USUV also affects the respiratory system and IHC of recent studies revealed the presence of USUV antigens in the lung [17,56]. A high number of findings in USUV positive birds were consistent with granulomatous pneumonia (43%; n=13/30). In birds, granulomatous pneumonia is often associated with fungal colonization of lungs and air sacs, which might be facilitated by immunosuppression [63].

An important aspect of this work was to partially characterize the viral strains circulating in Switzerland and ideally tracing a preliminary phylogeographic map of USUV in the country. USUV occurrence in Switzerland reflects the overall situation in Europe, especially in neighboring countries [64] with a predominance of European lineage and only one report of an African lineage. USUV (lineage Europe 1), in continental Europe was first detected in Italy in 1996 [9]. More recently, the variety of strains occurring in Europe has dramatically expanded with numerous new strains belonging to seven of the eight lineages known to date, including Europe 1-5, and Africa 2 and 3 [5,20,24]. The introduction of USUV to Europe has been regularly occurring from Africa during the last decades. Genetic variation of European lineages is predominantly the product of *in situ* evolution [65]. Differently, the African lineages appear to have been driven by extensive gene flow [65]. Starting in 2015 and with a major peak in 2018, USUV related bird mortality was observed in several European countries, including Belgium, France, Germany and the Netherlands [5,20,47] paralleled by an increase of clinical USUV cases and development of USUV specific antibodies in people [31,32,37,66]. In Switzerland, after the first detection of USUV in 2006 [11] consistent with a Europe 1 strain, mortality observed in captive and wild birds due to USUV was relatively negligible until 2015, when cases were observed more consistently, including several strains from Europe 3 lineage, suggesting a new introduction, or a previous lack of detection. The year 2018, stands out as the year accounting for most USUV associated casualties since its emergence in Switzerland. The high number of deaths was associated with the demonstration for the first time in Switzerland of the most diverse viral population recorded to date, with several subvariants of the Europe lineages (mainly Europe lineage 3) and for the first time, strains from the Africa 3 lineage. These strains and lineages have been circulating in neighboring countries in the previous years, such as Germany, Belgium, Netherlands, France [5] and Italy [67]. Accordingly, the recent epidemic in Switzerland was associated with a circulation of a major number of viral strains, including lineages and variants not previously recorded across the Swiss territory, as part of a pan-European pandemic. Nationally, USUV was mainly detected in the Central-North to North-Eastern regions (Plateau and Jura) and just one cluster was detected in the South (Alps region), suggesting a predominant introduction of the virus form Northern areas, and to a minor extent from the Southern territories. Although this could simply reflect a bias of analyzed specimens (most animals were delivered from Northern areas), this might also reflect the migratory route of birds flying from Africa back to Europe [68] or the closeness to Italy which may play an important role in the spread of USUV to the North, including Switzerland [69].

Although *Culex pipiens* (Diptera: Culicidae) is considered the main competent vector for USUV, other ornithophilic arthropods occurring in Switzerland, such as the native mosquito *Culex torrentium* [70], could potentially play a (marginal) role in the transmission of USUV and WNV [71]. The role played by the most common occurring tick (*Ixodes ricinus*) [72] and two invasive *Aedes* spp. (Diptera: Culicidae) mosquitos (the Asian tiger mosquito*, Ae. albopictus* and the East Asian bush mosquito*, Ae. japonicus*) [73,74] is questionable until now Vector-borne diseases are highly influenced by environmental changes and human activities [75].

The obvious increase of cases in birds in 2018 could be intuitively associated with a relative abundance of the main vector, *Cx. pipiens* [3,76], possibly occurred within the same time-frame and potentially being driven by favorable environmental conditions. In particular, high temperatures seem to be critical, more than high humidity (high precipitation) [77] to boost the vector population. Whereas we do not know the mosquito abundance for the whole Switzerland, we do know that the late spring and early summer of 2018 was extraordinarily hot and dry which could have resulted in increased mosquito density.

Furthermore, invasive species like the Asian tiger mosquito (*Ae albopictus*), in which USUV has been detected as well, are constantly expanding their territories, including Southern Switzerland [78]. Since its first detection in 2003 the Asian tiger mosquito seems to have established in the Southern parts of Switzerland [78] and continues to spread along main road connections into the Northern parts of Switzerland [79–81].

A specific statement of the WHO, supports this hypothesis clarifying that the high temperatures and extended rainy spells, followed by dry weather occurred in 2018 were key to boosting the mosquito population [82]. Furthermore, the closely related flavivirus WNV, another arthropod-borne virus, vectored mainly by *Cx. pipiens* as well, has also shown a significant peak of infections in humans and horses especially in Italy, Greece and Croatia in 2018 [44,83], but also in common blackbirds, northern goshawks and great grey owls in Germany [84].

Curiously, both WNV and USUV are vectored by the same mosquito species, *Culex pipiens*, additionally supporting the occurrence of environmental conditions favoring the increase of *Cx. pipiens* population and as consequentially, the increase of the outbreaks caused by the vectored viruses. Interestingly, to the best of our knowledge, besides being closely associated arbovirus and therefore following similar ecology, West-Nile Virus has only recently been detected in vectors in Switzerland [85] but not associated with obviously detected clinical diseases in wildlife or domestic animals, yet. This is in contrast to multiple detections in Germany [25,46,84,86], France [87,88], or Italy [53,68,89]. However, vaccination of domestic animals (horses) against WNV plus control measurements of invasive mosquito species such as *Aedes* spp. also affects native species as *Culex* spp. might impact the eco-epidemiology of Flaviviruses and clinical occurrence of infection [79].

Accordingly, the peak of avian mortality in summer and autumn in 2018 in Switzerland was likely the result of a combination of events, with on one side, the hot spring and summer, possibly contributing to boost the mosquito population and the chance of viral spread. On the other side, the genetic variability of multiple strains possibly poses challenges to the immune system of the hosts coping with continuously changing antigens. Starting already in 2019, the number of observed cases dropped significantly (4 in 2019 and in 2020 vs 44 cases in 2018). This could suggest on one side that the exceptional features of 2018 were not sustained by similar environmental conditions later on. On the other side, the likely reduction of the susceptible population during the summer of 2018, might have provided a lower number of potential hosts in the following year. Finally, a certain level of herd immunity might have been acquired in 2018 by the surviving birds, partially shielding them from the virus and the associated disease in the following season, as suggested and observed in other studies [77,90]. Nevertheless, despite the role of vectors is crucial, overwintering of the virus has been demonstrated locally and should be considered in the eco-epidemiology of the disease [91,92]. Moreover, an endemization of the USUV has been reported in neighboring countries [87] and can be expected in future, in Switzerland too.

Further observation and long-pathology and virology data sets will be fundamental to clarify these aspects and thoroughly understand the disease ecology of USUV. In particular, complementing host-pathogen interaction with environmental data, would provide the necessary vantage point to understand *disease in the making*.

Finally, this study broadens the knowledge on the disease and species distribution of USUV in Europe, adding critical information related to a geographically strategic country such as Switzerland, located in the heart of the Continent, at the crossroads of many birds-migratory routes.

## Material & Methods

### Study area and animals

Included in this study were 67 free-ranging wild birds collected in Switzerland and delivered at the FIWI between August 2015 and December 2020, which were conclusively confirmed USUV positive by reverse transcription PCR (RT-PCR). Switzerland is a small (surface area of 41,285 km^2^) Alpine country in central Europe which can be divided into three main biogeographical regions: the Jura in the north-western portion, the central Plateau and the Alps in the southern half of the country.

### Pathology

Full necropsy was carried out on all 67 birds and full or partial tissues sets were collected from each of them according to the degree of autolysis of each individual and integrity of their carcasses. When autolysis was severely advanced no tissues were collected for histology. Collected tissues comprised tongue, esophagus, trachea, lungs, heart, liver, spleen, gizzard, small and large intestine, kidneys, gonads, adrenals, cloaca, and brain. Furthermore, any additional organ showing macroscopic changes was also sampled. Tissues were fixed in 10% buffered formalin, processed routinely, sectioned at 5-μm-thick and stained with hematoxylin and eosin (HE) according to the accredited protocol of the Institute of Animal Pathology, Vetsuisse faculty Bern. Special stains including Periodic-Acid-Schiff (PAS), Grocott, Brown and Brenn’s Gram, and Giemsa stains were used as appropriate according to the initial findings of each bird.

Gross and histological lesions were graded according to an arbitrarily developed scheme according to the criteria summarized in Tables 1 and 2, respectively.

### USUV RT-PCR Reverse Transcription Polymerase Chain Reaction

Total RNA was extracted from the brain or the spleen of all the examined animals using the RNeasy Mini Kit (Qiagen, Hombrechticon, Switzerland) and converted into complementary DNA (cDNA) using the M-MLV reverse transcriptase kit (Promega, Madison, Wisconsin, USA) according to the manufacturers’ instructions. The cDNA was used as template for an in-house-developed reverse transcription PCR (RT-PCR) reaction detecting the partial sequence of the RNA polymerase gene of USUV. The latter was carried out using primers USUTU-Pol-FW (5’-TGAAAGGAGAGTGCCACACATGC-3’) and USUTU-Pol-RV (5’-TGAATGGTGGCTCATCTCACGC-3’) (Microsynth, Belgach, Switzerland). A total of 0.5μl of a 100μM solution of each of forward and reverse primer were added to a reaction mix. The reaction mix comprised 3μl of 10x reaction buffer, 3.75μl of MgCl_2_ (25Mm), 0.4μl of a 10mM dNTPs mix, 0.2μl (1U) of FIREpol DNA polymerase (Solis biodyne, Lucerna Chem, Luzern, Switzerland), and distilled deionized water with a final volume of 30μl. The cycling reaction was carried out on an Applied Biosystems Thermocycler 9600 Fast (Applied Biosystems, Foster City, Canada). The cycling protocol consisted of an initial denaturation step at 95°C for 3 minutes followed by 35 cycles including a denaturation step at 95°C for 30 seconds, an annealing step at 55°C for 30 seconds, and an elongation step at 72°C for 60 seconds. An additional elongation at 72°C for 10 minutes was run to exhaust the polymerase. The obtained amplicon was afterwards resolved in a 1.5% agarose gel and examined under ultraviolet light.

### Pan-Flavivirus RT-PCR

Samples which tested positive for the presence of USUV RNA by the RT-PCR described above were then further tested with a modified, widely used pan-flavivirus PCR protocol (Becker et al. 2012), which was considered ideal to compare the Swiss USUV strains with those available in public databases. Briefly, a total of 0.5μl of a 100μM solution of each forward (mFU1; 5’-TACAACATGATGGGAAAGCGAGAGAAAAA-3’) (Microsynth, Belgach, Switzerland) and reverse primer (cFD2; 5’-GTGTCCCAGCCGGCGGTGTCATCAGC-3’) (Microsynth, Belgach, Switzerland) were added to the reaction mix comprised 3μl of 10x reaction buffer, 3.75μl of MgCl_2_ (25Mm), 0.4μl of a 10mM dNTPs mix, 0.2μl (1U) of FIREpol DNA polymerase (Solis biodyne, Lucerna Chem, Luzern, Switzerland), and distilled deionized water up to a final volume of 30μl. The cycling reaction was carried out as described above with the annealing temperature of 57°C.

### Sequencing, assembly and phylogenetic analysis

The amplicons of each positive sample were submitted for automated Sanger sequencing for both strands using the same primers used for their amplification to a private biotech company (Microsynth SA, Belgach, Switzerland). Sequences were assessed for completeness and similarities using the software BLAST (https://blast.ncbi.nlm.nih.gov/Blast.cgi). All nucleotide ambiguities were checked manually directly on the chromatograms using the ABI PRISM ® software (Thermo Fischer Scientific^TM^, Basel, Switzerland).

Phylogenetic analysis was based on the partial sequence of the NS5 gene, which has been shown to provide a reliable proxy for the whole genome (Cadar et al. 2017a; Engel et al. 2016). The nucleotide sequences obtained from each sample were used for the phylogenetic analysis together with reference sequences from the different known USUV lineages, including Africa 1 (AY453412.1, KC754958.1), Africa 2 (KC754954.1), Africa 3 (KC754955.1, KC754956.1, KC754957.1, KM659877.1, MN122249.1), Europe 1 (NC_006551.1, JQ219843.1, EF206350.1), Europe 2 (HM569263.1, JF266698.1), Europe 3 (HM138716.1, HE599647.1, KJ859682.1, KJ859683.1, KY128481.1, KY263624.1, KY263623.1, KY263622.1, HM138716.1) Europe 4 (HM138718.1, JF834547.1) and Europe 5 (KY113091.1, KY113101.1) and a sequence of West Nile virus (MF797870.1) which was used as outgroup. Briefly, the sequences were aligned using Muscle (https://www.ebi.ac.uk/Tools/msa/muscle/) with standard settings. A maximum likelihood phylogenetic tree was built using Mega7 (Tamura et al. 2011) with standard settings, including 1,000 bootstraps replications, Tamura-Nei substitution model, uniform rates and nearest neighbor interchange (NNI).

### Data management and Statistics

Data management and descriptive statistics were conducted in Microsoft Excel 2016 (Microsoft Corporation, Redmond, Washington, USA). Statistical analyses were performed with R version 3.4.3 (http://www.r-project.org). Maps were designed with qGIS version 2.18.16 Las Palmas (Free Software Foundation Inc., Boston, USA, (http://www.qgis.org)) using standard settings.

### Environmental conditions

Environmental conditions recorded for the years 2015-2020 with particular attention for 2018 were obtained by the seasonal climate bulletin of the Swiss meteorological website [50,51], closely analyzing those conditions, which clearly differentiated 2018 from the other years of the investigated time window.

## Acknowledgements

We would like to thank all people that submitted diseases or dead bird in the framework of national surveillance program for wildlife health. And special thanks to the diagnostic assistants of the Wildlife Diagnostic Units for performing necropsy, namely (in alphabetical order) Ezgi Akdesir, Stéphanie Borel, Giuseppina Gelormini, Chloé Haas, Nelson Marreros, Iris Andrea Marti, Gaia Moore-Jones, Patrick Scherrer and Ursula Teubenbacher. Furthermore, we would like to thank Eliane Jemmi from FIWI for her help in analyzing the sequences of USUV strains and the respective nucleotide substitutions and Eleonora Flacio from the University of Applied Sciences and Arts of Southern Switzerland for her insights on mosquito ecology in Switzerland.

## References

1. Gould EA, Solomon T. Pathogenic flaviviruses. The Lancet. 2008; 371:500–9. doi: 10.1016/S0140-6736(08)60238-X.

2. Ashraf U, Ye J, Ruan X, Wan S, Zhu B, Cao S. Usutu virus: an emerging flavivirus in Europe. Viruses. 2015; 7:219–38. doi: 10.3390/v7010219 PMID: 25606971.

3. Nikolay B. A review of West Nile and Usutu virus co-circulation in Europe: how much do transmission cycles overlap. Trans R Soc Trop Med Hyg. 2015; 109:609–18. doi: 10.1093/trstmh/trv066 PMID: 26286946.

4. Roesch F, Fajardo A, Moratorio G, Vignuzzi M. Usutu Virus: An Arbovirus on the Rise. Viruses. 2019; 11. doi: 10.3390/v11070640 PMID: 31336826.

5. Cadar D, Lühken R, van der Jeugd H, Garigliany M, Ziegler U, Keller M, et al. Widespread activity of multiple lineages of Usutu virus, western Europe, 2016. Euro Surveill. 2017; 22. doi: 10.2807/1560-7917.ES.2017.22.4.30452 PMID: 28181903.

6. Calzolari M, Bonilauri P, Bellini R, Albieri A, Defilippo F, Maioli G, et al. Evidence of simultaneous circulation of West Nile and Usutu viruses in mosquitoes sampled in Emilia-Romagna region (Italy) in 2009. PLoS ONE. 2010; 5:e14324. doi: 10.1371/journal.pone.0014324 PMID: 21179462.

7. Williams MC, Simpson DI, Haddow A. J., Knight EM. The Isolation of West Nile Virus from Man and of Usutu Virus from the Bird-Biting Mosquito Mansonia Aurites (Theobald) in the Entebbe Area of Uganda. Ann Trop Med Parasitol. 1964; 58:367–74. doi: 10.1080/00034983.1964.11686258 PMID: 14212897.

8. Weissenböck H, Kolodziejek J, Fragner K, Kuhn R, Pfeffer M, Nowotny N. Usutu virus activity in Austria, 2001–2002. Microbes Infect. 2003; 5:1132–6. doi: 10.1016/S1286-4579(03)00204-1.

9. Weissenböck H, Bakonyi T, Rossi G, Mani P, Nowotny N. Usutu virus, Italy, 1996. Emerging Infect Dis. 2013; 19:274–7. doi: 10.3201/eid1902.121191 PMID: 23347844.

10. Bakonyi T, Erdélyi K, Ursu K, Ferenczi E, Csörgo T, Lussy H, et al. Emergence of Usutu virus in Hungary. J Clin Microbiol. 2007; 45:3870–4. doi: 10.1128/JCM.01390-07 PMID: 17913929.

11. Steinmetz HW, Bakonyi T, Weissenböck H, Hatt J-M, Eulenberger U, Robert N, et al. Emergence and establishment of Usutu virus infection in wild and captive avian species in and around Zurich, Switzerland—genomic and pathologic comparison to other central European outbreaks. Vet Microbiol. 2011; 148:207–12. doi: 10.1016/j.vetmic.2010.09.018 PMID: 20980109.

12. Bakonyi T, Busquets N, Nowotny N. Comparison of complete genome sequences of Usutu virus strains detected in Spain, Central Europe, and Africa. Vector Borne Zoonotic Dis. 2014; 14:324–9. doi: 10.1089/vbz.2013.1510 PMID: 24746182.

13. Savini G, Monaco F, Terregino C, Di Gennaro A, Bano L, Pinoni C, et al. Usutu virus in Italy: an emergence or a silent infection. Vet Microbiol. 2011; 151:264–74. doi: 10.1016/j.vetmic.2011.03.036 PMID: 21550731.

14. Jöst H, Bialonski A, Maus D, Sambri V, Eiden M, Groschup MH, et al. Isolation of usutu virus in Germany. Am J Trop Med Hyg. 2011; 85:551–3. doi: 10.4269/ajtmh.2011.11-0248 PMID: 21896821.

15. Hubálek Z, Rudolf I, Čapek M, Bakonyi T, Betášová L, Nowotny N. Usutu virus in blackbirds (Turdus merula), Czech Republic, 2011-2012. Transbound Emerg Dis. 2014; 61:273–6. Epub 2012/10/24. doi: 10.1111/tbed.12025 PMID: 23095331.

16. Garigliany M, Linden A, Gilliau G, Levy E, Sarlet M, Franssen M, et al. Usutu virus, Belgium, 2016. Infect Genet Evol. 2017; 48:116–9. doi: 10.1016/j.meegid.2016.12.023 PMID: 28017913.

17. Benzarti E, Sarlet M, Franssen M, Cadar D, Schmidt-Chanasit J, Rivas JF, et al. Usutu Virus Epizootic in Belgium in 2017 and 2018: Evidence of Virus Endemization and Ongoing Introduction Events. Vector Borne Zoonotic Dis. 2020; 20:43–50. doi: 10.1089/vbz.2019.2469 PMID: 31479400.

18. Rouffaer LO, Steensels M, Verlinden M, Vervaeke M, Boonyarittichaikij R, Martel A, et al. Usutu Virus Epizootic and Plasmodium Coinfection in Eurasian Blackbirds (*Turdus merula*) in Flanders, Belgium. J Wildl Dis. 2018; 54:859–62. doi: 10.7589/2017-07-163 PMID: 29889004.

19. Lecollinet S, Blanchard Y, Manson C, Lowenski S, Laloy E, Quenault H, et al. Dual Emergence of Usutu Virus in Common Blackbirds, Eastern France, 2015. Emerging Infect Dis. 2016; 22:2225–7. doi: 10.3201/eid2212.161272 PMID: 27869608.

20. Rijks JM, Kik ML, Slaterus R, Foppen R, Stroo A, IJzer J, et al. Widespread Usutu virus outbreak in birds in the Netherlands, 2016. Euro Surveill. 2016; 21. doi: 10.2807/1560-7917.ES.2016.21.45.30391 PMID: 27918257.

21. Folly AJ, Lawson B, Lean FZ, McCracken F, Spiro S, John SK, et al. Detection of Usutu virus infection in wild birds in the United Kingdom, 2020. Euro Surveill. 2020; 25. doi: 10.2807/1560-7917.ES.2020.25.41.2001732 PMID: 33063656.

22. Snoeck CJ, Sausy A, Losch S, Wildschutz F, Bourg M, Hübschen JM. Usutu Virus Africa 3 Lineage, Luxembourg, 2020. Emerging Infect Dis. 2022; 28:1076–9. doi: 10.3201/eid2805.212012 PMID: 35447065.

23. Angeloni G, Bertola M, Lazzaro E, Morini M, Masi G, Sinigaglia A, et al. Epidemiology, surveillance and diagnosis of Usutu virus infection in the EU/EEA, 2012 to 2021. Euro Surveill. 2023; 28. doi: 10.2807/1560-7917.ES.2023.28.33.2200929 PMID: 37589592.

24. Benzarti E, Linden A, Desmecht D, Garigliany M. Mosquito-borne epornitic flaviviruses: an update and review. J Gen Virol. 2019; 100:119–32. doi: 10.1099/jgv.0.001203 PMID: 30628886.

25. Michel F, Fischer D, Eiden M, Fast C, Reuschel M, Müller K, et al. West Nile Virus and Usutu Virus Monitoring of Wild Birds in Germany. Int J Environ Res Public Health. 2018; 15. Epub 2018/01/22. doi: 10.3390/ijerph15010171 PMID: 29361762.

26. Cadar D, Becker N, Campos RdM, Börstler J, Jöst H, Schmidt-Chanasit J. Usutu virus in bats, Germany, 2013. Emerging Infect Dis. 2014; 20:1771–3. doi: 10.3201/eid2010.140909 PMID: 25271769.

27. Durand B, Haskouri H, Lowenski S, Vachiery N, Beck C, Lecollinet S. Seroprevalence of West Nile and Usutu viruses in military working horses and dogs, Morocco, 2012: dog as an alternative WNV sentinel species. Epidemiol Infect. 2016; 144:1857–64. doi: 10.1017/S095026881600011X PMID: 26838515.

28. Barbic L, Vilibic-Cavlek T, Listes E, Stevanovic V, Gjenero-Margan I, Ljubin-Sternak S, et al. Demonstration of Usutu virus antibodies in horses, Croatia. Vector Borne Zoonotic Dis. 2013; 13:772–4. doi: 10.1089/vbz.2012.1236 PMID: 23808977.

29. García-Bocanegra I, Paniagua J, Gutiérrez-Guzmán AV, Lecollinet S, Boadella M, Arenas-Montes A, et al. Spatio-temporal trends and risk factors affecting West Nile virus and related flavivirus exposure in Spanish wild ruminants. BMC Vet Res. 2016; 12:249. doi: 10.1186/s12917-016-0876-4 PMID: 27829427.

30. Allering L, Jöst A, Emmerich P, Günther S, Lattwein E, Schmidt M, et al. Detection of Usutu virus infection in a healthy blood donor from south-west Germany, 2012. Euro Surveill. 2017; 2012:20341. Available from: 10.2807/ese.17.50.20341-en.

31. Pacenti M, Sinigaglia A, Martello T, Rui ME de, Franchin E, Pagni S, et al. Clinical and virological findings in patients with Usutu virus infection, northern Italy, 2018. Euro Surveill. 2019; 24. doi: 10.2807/1560-7917.ES.2019.24.47.1900180 PMID: 31771697.

32. Simonin Y, Sillam O, Carles MJ, Gutierrez S, Gil P, Constant O, et al. Human Usutu Virus Infection with Atypical Neurologic Presentation, Montpellier, France, 2016. Emerging Infect Dis. 2018; 24:875–8. doi: 10.3201/eid2405.171122 PMID: 29664365.

33. Garigliany M-M, Marlier D, Tenner-Racz K, Eiden M, Cassart D, Gandar F, et al. Detection of Usutu virus in a bullfinch (*Pyrrhula pyrrhula*) and a great spotted woodpecker (*Dendrocopos major)* in north-west Europe. Vet J. 2014; 199:191–3. doi: 10.1016/j.tvjl.2013.10.017 PMID: 24268481.

34. Weissenböck H, Bakonyi T, Chvala S, Nowotny N. Experimental Usutu virus infection of suckling mice causes neuronal and glial cell apoptosis and demyelination. Acta Neuropathol. 2004; 108:453–60. doi: 10.1007/s00401-004-0916-1 PMID: 15372281.

35. Weissenböck H, Kolodziejek J, Url A, Lussy H, Rebel-Bauder B, Nowotny N. Emergence of Usutu virus, an African Mosquito-Borne Flavivirus of the Japanese Encephalitis Virus Group, Central Europe. Emerging Infect Dis. 2008; 2002:652–6.

36. Cadar D, Maier P, Müller S, Kress J, Chudy M, Bialonski A, et al. Blood donor screening for West Nile virus (WNV) revealed acute Usutu virus (USUV) infection, Germany, September 2016. Euro Surveill. 2017; 22. doi: 10.2807/1560-7917.ES.2017.22.14.30501 PMID: 28422005.

37. Cadar D, Simonin Y. Human Usutu Virus Infections in Europe: A New Risk on Horizon. Viruses. 2022; 15. Epub 2022/12/27. doi: 10.3390/v15010077 PMID: 36680117.

38. Gaibani P, Barp N, Massari M, Negri EA, Rossini G, Vocale C, et al. Case report of Usutu virus infection in an immunocompromised patient in Italy, 2022. J Neurovirol. 2023; 29:364–6. Epub 2023/05/25. doi: 10.1007/s13365-023-01148-w PMID: 37227671.

39. Pecorari M. First human case of Usutu virus neuroinvasive infection, Italy, August-September 2009. Euro Surveill. 2009; 14:19446. Available from: http://www.eurosurveillance.org/ViewArticle.aspx?ArticleId=19446.

40. Nikolay B, Diallo M, Boye CSB, Sall AA. Usutu virus in Africa. Vector Borne Zoonotic Dis. 2011; 11:1417–23. doi: 10.1089/vbz.2011.0631 PMID: 21767160.

41. Clé M, Beck C, Salinas S, Lecollinet S, Gutierrez S, van de Perre P, et al. Usutu virus: A new threat. Epidemiol Infect. 2019; 147:e232. doi: 10.1017/S0950268819001213 PMID: 31364580.

42. Ryser-Degiorgis M-P, Segner H. National competence center for wildlife diseases in Switzerland: Mandate, development and current strategies. Schweiz Arch Tierheilkd. 2015; 157:255–66. doi: 10.17236/sat00019 PMID: 26753341.

43. Weidinger P, Kolodziejek J, Bakonyi T, Brunthaler R, Erdélyi K, Weissenböck H, et al. Different dynamics of Usutu virus infections in Austria and Hungary, 2017-2018. Transbound Emerg Dis. 2020; 67:298–307. doi: 10.1111/tbed.13351 PMID: 31505099.

44. Vilibic-Cavlek T, Savic V, Petrovic T, Toplak I, Barbic L, Petric D, et al. Emerging Trends in the Epidemiology of West Nile and Usutu Virus Infections in Southern Europe. Front Vet Sci. 2019; 6:437. doi: 10.3389/fvets.2019.00437 PMID: 31867347.

45. Čabanová V, Šikutová S, Straková P, Šebesta O, Vichová B, Zubríková D, et al. Co-Circulation of West Nile and Usutu Flaviviruses in Mosquitoes in Slovakia, 2018. Viruses. 2019; 11. doi: 10.3390/v11070639 PMID: 31336825.

46. Michel F, Sieg M, Fischer D, Keller M, Eiden M, Reuschel M, et al. Evidence for West Nile Virus and Usutu Virus Infections in Wild and Resident Birds in Germany, 2017 and 2018. Viruses. 2019; 11. doi: 10.3390/v11070674 PMID: 31340516.

47. Oude Munnink BB, Münger E, Nieuwenhuijse DF, Kohl R, van der Linden A, Schapendonk CME, et al. Genomic monitoring to understand the emergence and spread of Usutu virus in the Netherlands, 2016-2018. Sci Rep. 2020; 10:2798. doi: 10.1038/s41598-020-59692-y PMID: 32071379.

48. Groen TA, L’Ambert G, Bellini R, Chaskopoulou A, Petric D, Zgomba M, et al. Ecology of West Nile virus across four European countries: empirical modelling of the *Culex pipiens* abundance dynamics as a function of weather. Parasit Vectors. 2017; 10:524. Epub 2017/10/26. doi: 10.1186/s13071-017-2484-y PMID: 29070056.

49. Pachka H, Annelise T, Alan K, Power T, Patrick K, Véronique C, et al. Rift Valley fever vector diversity and impact of meteorological and environmental factors on *Culex pipiens* dynamics in the Okavango Delta, Botswana. Parasit Vectors. 2016; 9:434. Epub 2016/08/08. doi: 10.1186/s13071-016-1712-1 PMID: 27502246.

50. Bundesamt für Meteorologie und Klimatologie MeteoSchweiz. Klimareport 2018.

51. Bundesamt für Meteorologie und Klimatologie MeteoSchweiz. Hitze und Trockenheit im Sommerhalbjahr 2018 – eine klimatologische Übersicht. Available from: https://www.meteoschweiz.admin.ch/home/service-und-publikationen/publikationen.html?topic=/content/meteoswiss/tags/topics/forschung-und-zusammenarbeit/publication/fachbeitraege&query=Hitze+und+Trockenheit+im+Sommerhalbjahr+2018&topic=/content/meteoswiss/tags/topics/klima&publicationYear=&pageIndex=0.

52. World Health Organization. West Nile virus infections spike in southern and central Europe. World Health Organization 2018 [updated 16 Jul 2020; cited 5 Aug 2020]. Available from: https://www.euro.who.int/en/countries/italy/news/news/2018/8/west-nile-virus-infections-spike-in-southern-and-central-europe.

53. Lauriano A, Rossi A, Galletti G, Casadei G, Santi A, Rubini S, et al. West Nile and Usutu Viruses’ Surveillance in Birds of the Province of Ferrara, Italy, from 2015 to 2019. Viruses. 2021; 13. Epub 2021/07/14. doi: 10.3390/v13071367 PMID: 34372573.

54. Chvala S, Kolodziejek J, Nowotny N, Weissenböck H. Pathology and viral distribution in fatal Usutu virus infections of birds from the 2001 and 2002 outbreaks in Austria. J Comp Pathol. 2004; 131:176–85. doi: 10.1016/j.jcpa.2004.03.004 PMID: 15276857.

55. Agliani G, Giglia G, Marshall EM, Gröne A, Rockx BHG, van den Brand JMA. Pathological features of West Nile and Usutu virus natural infections in wild and domestic animals and in humans: A comparative review. One Health. 2023; 16:100525. Epub 2023/03/10. doi: 10.1016/j.onehlt.2023.100525 PMID: 37363223.

56. Giglia G, Agliani G, Munnink BBO, Sikkema RS, Mandara MT, Lepri E, et al. Pathology and Pathogenesis of Eurasian Blackbirds (*Turdus merula*) Naturally Infected with Usutu Virus. Viruses. 2021; 13. Epub 2021/07/28. doi: 10.3390/v13081481 PMID: 34452347.

57. Störk T, Le Roi M de, Haverkamp A-K, Jesse ST, Peters M, Fast C, et al. Analysis of avian Usutu virus infections in Germany from 2011 to 2018 with focus on dsRNA detection to demonstrate viral infections. Sci Rep. 2021; 11:24191. Epub 2021/12/17. doi: 10.1038/s41598-021-03638-5 PMID: 34921222.

58. Ka MB, Olive D, Mege J-L. Modulation of monocyte subsets in infectious diseases. WJI. 2014; 4:185. doi: 10.5411/wji.v4.i3.185.

59. Nikitina E, Larionova I, Choinzonov E, Kzhyshkowska J. Monocytes and Macrophages as Viral Targets and Reservoirs. Int J Mol Sci. 2018; 19. Epub 2018/09/18. doi: 10.3390/ijms19092821 PMID: 30231586.

60. Rios M, Zhang MJ, Grinev A, Srinivasan K, Daniel S, Wood O, et al. Monocytes-macrophages are a potential target in human infection with West Nile virus through blood transfusion. Transfusion. 2006; 46:659–67. doi: 10.1111/j.1537-2995.2006.00769.x PMID: 16584445.

61. Zimmerman MG, Bowen JR, McDonald CE, Pulendran B, Suthar MS. West Nile Virus Infection Blocks Inflammatory Response and T Cell Costimulatory Capacity of Human Monocyte-Derived Dendritic Cells. J Virol. 2019; 93. Epub 2019/11/13. doi: 10.1128/JVI.00664-19 PMID: 31534040.

62. Origgi FC, Plattet P, Sattler U, Robert N, Casaubon J, Mavrot F, et al. Emergence of canine distemper virus strains with modified molecular signature and enhanced neuronal tropism leading to high mortality in wild carnivores. Vet Pathol. 2012; 49:913–29. doi: 10.1177/0300985812436743 PMID: 22362965.

63. Nemeth NM, Gonzalez-Astudillo V, Oesterle PT, Howerth EW. A 5-Year Retrospective Review of Avian Diseases Diagnosed at the Department of Pathology, University of Georgia. J Comp Pathol. 2016; 155:105–20. Epub 2016/06/18. doi: 10.1016/j.jcpa.2016.05.006 PMID: 27329003.

64. Vilibic-Cavlek T, Petrovic T, Savic V, Barbic L, Tabain I, Stevanovic V, et al. Epidemiology of Usutu Virus: The European Scenario. Pathogens. 2020; 9. Epub 2020/08/26. doi: 10.3390/pathogens9090699 PMID: 32858963.

65. Engel D, Jöst H, Wink M, Börstler J, Bosch S, Garigliany M-M, et al. Reconstruction of the Evolutionary History and Dispersal of Usutu Virus, a Neglected Emerging Arbovirus in Europe and Africa. mBio. 2016; 7:e01938–15. doi: 10.1128/mBio.01938-15 PMID: 26838717.

66. Zaaijer HL, Slot E, Molier M, Reusken CBEM, Koppelman MHGM. Usutu virus infection in Dutch blood donors. Transfusion. 2019; 59:2931–7. doi: 10.1111/trf.15444 PMID: 31270821.

67. Calzolari M, Chiapponi C, Bonilauri P, Lelli D, Baioni L, Barbieri I, et al. Co-circulation of two Usutu virus strains in Northern Italy between 2009 and 2014. Infect Genet Evol. 2017; 51:255–62. doi: 10.1016/j.meegid.2017.03.022 PMID: 28341546.

68. Mancuso E, Cecere JG, Iapaolo F, Di Gennaro A, Sacchi M, Savini G, et al. West Nile and Usutu Virus Introduction via Migratory Birds: A Retrospective Analysis in Italy. Viruses. 2022; 14. Epub 2022/02/17. doi: 10.3390/v14020416 PMID: 35216009.

69. Zecchin B, Fusaro A, Milani A, Schivo A, Ravagnan S, Ormelli S, et al. The central role of Italy in the spatial spread of USUTU virus in Europe. Virus Evol. 2021; 7:veab048. Epub 2021/08/31. doi: 10.1093/ve/veab048 PMID: 34513027.

70. Kubacki J, Hardmeier I, Qi W, Flacio E, Tonolla M, Fraefel C. Complete Genome Sequence of a Rhabdovirus Strain from Culex Mosquitos Collected in Southern Switzerland. Microbiol Resour Announc. 2021; 10. Epub 2021/01/07. doi: 10.1128/mra.01234-20 PMID: 33414339.

71. Fros JJ, Miesen P, Vogels CB, Gaibani P, Sambri V, Martina BE, et al. Comparative Usutu and West Nile virus transmission potential by local *Culex pipiens* mosquitoes in north-western Europe. One Health. 2015; 1:31–6. doi: 10.1016/j.onehlt.2015.08.002 PMID: 28616462.

72. Bakker JW, Münger E, Esser HJ, Sikkema RS, Boer WF de, Sprong H, et al. Ixodes ricinus as potential vector for Usutu virus. PLoS Negl Trop Dis. 2024; 18:e0012172. Epub 2024/07/10. doi: 10.1371/journal.pntd.0012172 PMID: 38985837.

73. Puggioli A, Bonilauri P, Calzolari M, Lelli D, Carrieri M, Urbanelli S, et al. Does *Aedes albopictus* (Diptera: Culicidae) play any role in Usutu virus transmission in Northern Italy? Experimental oral infection and field evidences. Acta Trop. 2017; 172:192–6. doi: 10.1016/j.actatropica.2017.05.006 PMID: 28495404.

74. Linthout C, Martins AD, Wit M de, Delecroix C, Abbo SR, Pijlman GP, et al. The potential role of the Asian bush mosquito *Aedes japonicus* as spillover vector for West Nile virus in the Netherlands. Parasit Vectors. 2024; 17:262. Epub 2024/06/17. doi: 10.1186/s13071-024-06279-5 PMID: 38886805.

75. Souza WM de, Weaver SC. Effects of climate change and human activities on vector-borne diseases. Nat Rev Microbiol. 2024; 22:476–91. Epub 2024/03/14. doi: 10.1038/s41579-024-01026-0 PMID: 38486116.

76. Becker N, Jöst H, Ziegler U, Eiden M, Höper D, Emmerich P, et al. Epizootic emergence of Usutu virus in wild and captive birds in Germany. PLoS ONE. 2012; 7:e32604. doi: 10.1371/journal.pone.0032604 PMID: 22389712.

77. Rubel F, Brugger K, Hantel M, Chvala-Mannsberger S, Bakonyi T, Weissenböck H, et al. Explaining Usutu virus dynamics in Austria: model development and calibration. Prev Vet Med. 2008; 85:166–86. doi: 10.1016/j.prevetmed.2008.01.006 PMID: 18314208.

78. Flacio E, Engeler L, Tonolla M, Müller P. Spread and establishment of *Aedes albopictus* in southern Switzerland between 2003 and 2014: an analysis of oviposition data and weather conditions. Parasit Vectors. 2016; 9:304. Epub 2016/05/26. doi: 10.1186/s13071-016-1577-3 PMID: 27229686.

79. Ravasi D, Mangili F, Huber D, Cannata M, Strigaro D, Flacio E. The effects of microclimatic winter conditions in urban areas on the risk of establishment for *Aedes albopictus*. Sci Rep. 2022; 12:15967. Epub 2022/09/24. doi: 10.1038/s41598-022-20436-9 PMID: 36153403.

80. Müller P, Engeler L, Vavassori L, Suter T, Guidi V, Gschwind M, et al. Surveillance of invasive *Aedes* mosquitoes along Swiss traffic axes reveals different dispersal modes for Aedes albopictus and Ae. japonicus. PLoS Negl Trop Dis. 2020; 14:e0008705. Epub 2020/09/28. doi: 10.1371/journal.pntd.0008705 PMID: 32986704.

81. Infofauna. info fauna carto [updated 7 Oct 2024; cited 29 Apr 2025]. Available from: https://lepus.infofauna.ch/carto/35972.

82. ALDERMAN DJ. Geographical spread of bacterial and fungal diseases of crustaceans. Revue Scientifique et Technique de l’OIE. 1996; 15:603–32. doi: 10.20506/rst.15.2.943 PMID: 8890383.

83. European Centre for Disease Prevention and Control (ECDC). Historical data by year - West Nile fever seasonal surveillance [updated 5 Dec 2019; cited 26 Jun 2020]. Available from: https://www.ecdc.europa.eu/en/west-nile-fever/surveillance-and-disease-data/historical.

84. Ziegler U, Lühken R, Keller M, Cadar D, van der Grinten E, Michel F, et al. West Nile virus epizootic in Germany, 2018. Antiviral Res. 2019; 162:39–43. doi: 10.1016/j.antiviral.2018.12.005 PMID: 30550796.

85. Cazzin S, Liechti N, Jandrasits D, Flacio E, Beuret C, Engler O, et al. First Detection of West Nile Virus Lineage 2 in Mosquitoes in Switzerland, 2022. Pathogens. 2023; 12. Epub 2023/12/07. doi: 10.3390/pathogens12121424 PMID: 38133307.

86. Santos PD, Michel F, Wylezich C, Höper D, Keller M, Holicki CM, et al. Co-infections: Simultaneous detections of West Nile virus and Usutu virus in birds from Germany. Transbound Emerg Dis. 2022; 69:776–92. Epub 2021/03/31. doi: 10.1111/tbed.14050 PMID: 33655706.

87. Constant O, Gil P, Barthelemy J, Bolloré K, Foulongne V, Desmetz C, et al. One Health surveillance of West Nile and Usutu viruses: a repeated cross-sectional study exploring seroprevalence and endemicity in Southern France, 2016 to 2020. Euro Surveill. 2022; 27. doi: 10.2807/1560-7917.ES.2022.27.25.2200068 PMID: 35748300.

88. Beck C, Leparc Goffart I, Franke F, Gonzalez G, Dumarest M, Lowenski S, et al. Contrasted Epidemiological Patterns of West Nile Virus Lineages 1 and 2 Infections in France from 2015 to 2019. Pathogens. 2020; 9. Epub 2020/10/30. doi: 10.3390/pathogens9110908 PMID: 33143300.

89. Calzolari M, Gaibani P, Bellini R, Defilippo F, Pierro A, Albieri A, et al. Mosquito, bird and human surveillance of West Nile and Usutu viruses in Emilia-Romagna Region (Italy) in 2010. PLoS ONE. 2012; 7:e38058. doi: 10.1371/journal.pone.0038058 PMID: 22666446.

90. Meister T, Lussy H, Bakonyi T, Sikutová S, Rudolf I, Vogl W, et al. Serological evidence of continuing high Usutu virus (*Flaviviridae*) activity and establishment of herd immunity in wild birds in Austria. Vet Microbiol. 2008; 127:237–48. doi: 10.1016/j.vetmic.2007.08.023 PMID: 17869454.

91. Folly AJ, Sewgobind S, Hernández-Triana LM, Mansfield KL, Lean FZX, Lawson B, et al. Evidence for overwintering and autochthonous transmission of Usutu virus to wild birds following its redetection in the United Kingdom. Transbound Emerg Dis. 2022; 69:3684–92. Epub 2022/10/25. doi: 10.1111/tbed.14738 PMID: 36217722.

92. Blom R, Schrama MJJ, Spitzen J, Weller BFM, van der Linden A, Sikkema RS, et al. Arbovirus persistence in North-Western Europe: Are mosquitoes the only overwintering pathway. One Health. 2023; 16:100467. Epub 2022/12/01. doi: 10.1016/j.onehlt.2022.100467 PMID: 36531660.

